# Identification and quantification of transposable element transcripts using Long-Read RNA-seq in *Drosophila* germline tissues

**DOI:** 10.1101/2023.05.27.542554

**Authors:** Rita Rebollo, Pierre Gerenton, Eric Cumunel, Arnaud Mary, François Sabot, Nelly Burlet, Benjamin Gillet, Sandrine Hughes, Daniel S. Oliveira, Clément Goubert, Marie Fablet, Cristina Vieira, Vincent Lacroix

## Abstract

Transposable elements (TEs) are repeated DNA sequences potentially able to move throughout the genome. In addition to their inherent mutagenic effects, TEs can disrupt nearby genes by donating their intrinsic regulatory sequences, for instance, promoting the ectopic expression of a cellular gene. TE transcription is therefore not only necessary for TE transposition *per se* but can also be associated with TE-gene fusion transcripts, and in some cases, be the product of pervasive transcription. Hence, correctly determining the transcription state of a TE copy is essential to apprehend the impact of the TE in the host genome. Methods to identify and quantify TE transcription have mostly relied on short RNA-seq reads to estimate TE expression at the family level while using specific algorithms to discriminate copy-specific transcription. However, assigning short reads to their correct genomic location, and genomic feature is not trivial. Here we retrieved full-length cDNA (TeloPrime, Lexogen) of *Drosophila melanogaster* gonads and sequenced them using Oxford Nanopore Technologies. We show that long-read RNA-seq can be used to identify and quantify transcribed TEs at the copy level. In particular, TE insertions overlapping annotated genes are better estimated using long reads than short reads. Nevertheless, long TE transcripts (> 4.5 kb) are not well captured. Most expressed TE insertions correspond to copies that have lost their ability to transpose, and within a family, only a few copies are indeed expressed. Long-read sequencing also allowed the identification of spliced transcripts for around 107 TE copies. Overall, this first comparison of TEs between testes and ovaries uncovers differences in their transcriptional landscape, at the subclass and insertion level.

## Introduction

Transposable elements (TEs) are widespread DNA sequences that can move around genomes in a process called transposition (Bourque et al., 2018). TEs can transpose either using an RNA intermediate, in a copy-and-paste mechanism, *i.e.* retrotransposons, or directly through a DNA molecule using different cut-and-paste strategies, *i.e.* DNA transposons. In both cases, the synthesis of a messenger RNA is a fundamental step allowing the production of the transposition machinery, and hence promoting TE replication in the host genome. TE transposition is *per se* a mutational process, and several host mechanisms are in place in order to avoid novel TE insertions, including chromatin remodelling factors, DNA methylation, and small RNAs (Slotkin & Martienssen, 2007). For instance, in *Drosophila melanogaster* ovaries, TEs are the target of piwi-interacting RNAs (piRNAs) that promote TE transcript cleavage, but also deposition of repressive chromatin marks within the TE insertion, blocking any further transcription (Fabry et al., 2021).

In order to appreciate the dynamics of TE regulation, an accurate measure of TE expression is required, including copy-specific information (Lanciano & Cristofari, 2020). While such analyses may be easily obtained in genomes composed of mostly ancient TE copies, discrimination of young TE families, such as LINE-1, AluY and SVA in humans, or the study of genomes composed of mostly active copies as seen in many insects, remains a complex feat. Indeed, TE copies belonging to the same TE family have, by definition (Wicker et al., 2007), more than 80 % of sequence identity, hampering the study of TE regulation and consequently TE expression in a copy-specific manner (Lanciano & Cristofari, 2020). Most genome-wide analyses interested in TE expression, and even their regulation, focus on TE family-level analysis, where short reads are mapped either against TE consensus sequences or to the genome/TE copy sequences followed by grouping of read counts at the family level (TEcount from the TEtools package (Lerat et al., 2017), TEtranscripts (Jin et al., 2015)). In the past years, many methods have surfaced to take advantage of short-read sequencing datasets and circumvent the multi-mapping problem in order to develop copy-level analysis (for a review see (Lanciano & Cristofari, 2020)). These methods are based on different algorithms that can statistically reassign multi-mapped reads to unique locations, for instance with the expectation-maximization algorithm used in TEtranscripts (Jin et al., 2015), SQuIRE (Yang et al., 2019) and Telescope (Bendall et al., 2019).

In the past years, long-read sequencing has become an attractive alternative to study TE biology. Such reads are able to refine TE annotation (Jiang et al., 2019; Panda & Slotkin, 2020), pinpoint new TE insertions (Mohamed et al., 2020; Rech et al., 2022), determine TE DNA methylation rates at the copy level (Ewing et al., 2020), estimate TE expression (Berrens et al., 2022), and finally, detect TE-gene fusion transcripts (Panda & Slotkin, 2020; Dai et al., 2021; Babarinde et al., 2021). Furthermore, long-read RNA sequencing can not only determine which TE copies are expressed but also discriminate between isoforms of a single TE copy produced by alternative splicing. Indeed, TE alternative transcripts have been described in the very first studies of TEs, using techniques such as northern blot (Belancio et al., 2006), but concomitantly with accessible short-read genome-wide analysis, low interest has been given to TE transcript integrity. Nonetheless, TE isoforms have been shown to participate in TE regulation, as observed for the P element in *D. melanogaster*, where a specific germline isoform encodes a functional transposase protein, while in somatic tissues, another isoform acts as a P element repressor (Laski et al., 1986). The regulation of such tissue-specific splicing has recently been attributed to piRNA-directed heterochromatinization of P element copies (Teixeira et al., 2017). The retrotransposon *Gypsy* also produces two isoforms, including an envelope-encoding infectious germline isoform, also controlled by piRNA-guided repressive chromatin marks (Pélisson et al., 1994; Teixeira et al., 2017). Recently, Panda and Slotkin produced long-read RNA sequencing of *Arabidopsis thaliana* lines with defects in TE regulatory mechanisms (Panda & Slotkin, 2020), and were able to annotate TE transcripts, pinpoint TE splicing isoforms, and most importantly, demonstrate that properly spliced TE transcripts are protected from small RNA degradation.

*D. melanogaster* harbours around 12-20% of TE content, and recent studies have suggested that 24 TE superfamilies are potentially active (Adrion et al., 2017). Nevertheless, no indication of which copies are active has been documented. Here, we describe a bioinformatics procedure using long-read RNA sequencing, which enables the efficient identification of TE-expressed loci and variation in TE transcript structure and splicing. Furthermore, our procedure is powerful enough to uncover tissue-specific differences, as illustrated by comparing testes and ovaries data.

## Methods

### Reference genome and annotation

The dmgoth101 genome assembly was produced from Oxford Nanopore Technologies (ONT) long-read DNA sequencing and described in (Mohamed et al., 2020). Genome assembly has been deposited in the European Nucleotide Archive (ENA) under accession number PRJEB50024, assembly GCA_927717585.1. Gene annotation was performed as described in (Fablet et al., 2023). Briefly, gene annotation files were retrieved from Flybase (dmel-all-r6.46.gtf.gz) along with the matching genome sequence (fasta/dmel-all-chromosome-r6.46.fasta.gz). We then used LiftOff v1.6.1 (Shumate & Salzberg, 2021) with the command liftoff -g dmel-all-r6.46.gtf -f feature_types.txt -o dmgoth101.txt -u unmapped_dmgoth101.txt -dir annotations -flank 0.2 dmgoth101_assembl_chrom.fasta dmel-all-chromosome-r6.46.fasta to lift over gene annotations from the references to the GCA_927717585.1 genome assembly. One should note that feature_types.txt is a two line txt file containing ‘gene’ and ‘exon’. In order to locate and count the reads aligned against TE insertions, we produced a GTF file with the position of each TE insertion. We have used RepeatMasker against the reference genome using the manually curated TE library produced by (Rech et al., 2022) and the parameters: -a (produce alignments with Kimura two-parameters divergence) -s (sensitive mode), -cutoff 200 (minimum Smith-Waterman score) and -no_is (do not search for bacterial insertion sequences). We then used RepeatCraft (Wong & Simakov, 2019) to merge overlapping TE annotations with the following command line repeatcraft.py -r GCA_927717585.1.contig_named.fasta.out.gff -u GCA_927717585.1.contig_named.fasta.out -c repeatcraft.cgf -o dm101-repeatcraft. Visualization of alignments of TE copies to their consensus sequences were performed using blastn (Altschul et al., 1990) with the consensus sequences from (Rech et al., 2022) that can also be found in the https://gitlab.inria.fr/erable/te_long_read/.

### Drosophila *rearing*

*D. melanogaster* dmgoth101 strain was previously described by (Mohamed et al., 2020). Briefly, an isofemale line was derived from a wild-type female *D. melanogaster* from Gotheron, France, sampled in 2014, and sib-mated for 30 generations. Flies were maintained in 12-hour light cycles, and 24° C, in vials with nutritive medium, in small-mass cultures with approximately 50 pairs at each generation.

### Long-read RNA-seq and analysis

#### RNA extraction and library construction

Forty-five pairs of ovaries and 62 pairs of testes were dissected in cold PBS 1X from 4 to 8-day-old adults. Total RNA was extracted using the QiagenRNeasy Plus Kit (Qiagen, reference 74104) after homogenization (using a pellet pestle motor) of the tissues. DNA contamination was controlled and removed by DNAse treatment for 30 minutes at 37°C (Ambion). Total RNA was visualized in agarose gel to check DNA contamination and RNA integrity before storing at -80°C. The two RNA extracts were quantified with RNA BR reagents on Qubit 4.0 (Thermo Fisher Scientific) and qualified with RNA ScreenTape on Tapestation 4150 instrument (Agilent Technologies), the results showing no limited quantity and a high quality of the RNA molecules (RIN >9.8). We then took advantage of the TeloPrime Full-Length cDNA Amplification kit V2 (Lexogen) in order to enrich ovary and testis total RNA in full-length cDNAs (Figure S1). One should note that the amplified cDNAs are smaller than ∼3.5 kb. This protocol is highly selective for mRNAs that are both capped and polyadenylated and allows their amplification. TeloPrime recommends 2 µg total RNA per reaction and we performed two reactions for testis (total of 4 µg) and three reactions for ovaries (total of 6 µg). We determined the optimal PCR cycle number for each sample by quantitative PCR. The quantity and quality of the cDNA produced were checked again with Qubit (dsDNA BR) and Tapestation (D5000 DNA ScreenTape) to confirm the correct amplification of the cDNA and absence of degradation in cDNA fragment length profiles. It is important to note that we do not have replicates for the long-read dataset as the primary goal for this experiment was to evaluate the potential of this technique to identify the largest number of expressed TE copies and isoforms. Enriched full-length cDNAs generated from ovaries and testes were then processed into libraries using the SQK-LSK109 ligation kit (ONT) using 3 µg as starting material. The two libraries were sequenced separately in two flow cells R10 (FLO-MIN110) with a MinION device (Guppy version 2.3.6 and fast basecalling). We obtained 1,236,000 reads for ovaries and 2,925,554 for testes that passed the default quality filter (>Q7). Data are available online at the BioProject PRJNA956863.

#### Mapping

The analysis performed here can be replicated through https://gitlab.inria.fr/erable/te_long_read/, a GitLab containing all the scripts along with links and/or methods to retrieve the datasets used. Quality control was performed with NanoPlot v1.41.6 (De Coster et al., 2018). The median read length was 1.18 kb for ovaries and 1.44 kb for testes, the N50 read length was 1.7 kb for ovaries and 2.19 kb for testes, and the median quality was 7.7 for ovaries and 8.4 for testes (Table S1, Figure S2). Reads were mapped to the dmgoth101 genome using minimap2 (version 2.26) (Li, 2018) with the splice preset parameter (exact command line given in the GitLab). Most of reads (91.3% for ovaries, 98.8% for testes) could be mapped to the genome (Table S1). Out of those mapped reads, the majority (98.8% for ovaries and 95.1% for testes) mapped to a unique location (*i.e.* had no secondary alignment), and the vast majority (99.9% for ovaries and 97.7% for testes) mapped to a unique best location (*i.e.* in presence of secondary alignments, one alignment has a score strictly higher than the others). Indeed, if a read has several alignments with the same alignment score, then this means the read stems from exact repeats in the genome and they cannot be told apart, hence, one cannot know which copy is transcribed. However, if a read has several alignments with distinct alignment scores, then it means that the read stems from inexact repeats. The presence of this read in the dataset means that one of the copies is transcribed and we consider that it is the one with the highest alignment score. While it could be possible that the read actually stems from the copy with suboptimal alignment, this is highly unlikely because it would mean that there is a sequencing error at the position of the divergence between the two copies of the repeat. A sequencing error in any other position of the read would cause a decrease in the alignment score of both locations. An example of a read that maps to several locations, one with an alignment score larger than the others is given in Figure S3.

#### Chimeric reads

We also noticed that some reads were only partially mapped to the genome. In practice the query coverage distribution is bimodal (Figure S4), 80% of reads have a query coverage centered on 90%, while the remaining 20% have a query coverage centered at 50%. A thorough inspection of the unmapped regions of these partially mapped reads reveals that they stem from transcripts located elsewhere on the genome. Given that the transcripts covered by the read are themselves fully covered (both the primary locus and the secondary locus), we think that these chimeras are artifactual and were probably generated during ligation steps as previously described (White et al., 2017). Here, we chose to focus on the locus corresponding to the primary alignment and discard the secondary loci. In practice, this corresponds to the longest of the two transcripts. We also ran the same analyses after completely discarding those 20% chimeric reads, but the quantification of TEs is essentially the same (R=0.992, Figure S5). In particular, no TE quantification is particularly affected by the inclusion/exclusion of chimeric reads. In order to further help users identify problems related to chimeric reads, we now also provide an additional column in Table S3 and Table S4 indicating, for each TE copy, the average number of soft-clipped bases. A particularly high value could be an indication that the chimeric reads are not associated to a library preparation issue, but to a structural variation absent from the reference genome. In our dataset, we find that the percentage of soft-clipped bases is similar for all TE copies (∼18%).

##### Feature assignment

Once a read is assigned to a genomic location, it does not yet mean that it is assigned to a genomic feature. In order to decide which reads could correspond to a TE, we applied the following filters. First, we selected all reads where the mapping location overlaps the annotation of a TE for at least one base. Then, we discarded all reads that covered less than 10 % of the annotated TE. On table S3 and S4, the percentage of TEs covered by a uniquely mapping read is depicted as “Mean_TE_Span” and explained on Figure S6. It is important to note that no filter is based on the number of basepairs or proportion of the read that extends beyond the TE boundaries, but this metric is present as “Mean_Bases_Outside_TE_Annotation”, and “Mean_percent_ofbase_inside_TE” on Tables S3 and S4 allowing for different analyses to be performed (Figure S6). Finally, in the case where a read mapped to a genomic location where there are several annotated features (a TE and a gene, or two TEs), we assign the read to the feature whose genomic interval (excluding introns) has the smallest symmetric difference with the one of the read. The rationale for introducing this filter is best explained with examples. Figure S7 corresponds to a TE annotation overlapping a gene annotation. All reads map to both features, but the gene is fully covered while the TE is only partially covered. We conclude that the gene is expressed, not the TE. Figure S8 corresponds to a genomic location where a TE insertion (DNAREP1_INE-1$2R_RaGOO$4901615$4901964) is located within the intron of a gene (*Gp210*). Some reads map to the gene and not the TE. Some reads map to both the TE and the gene. We assign these reads to the TE because the coverage of symmetric difference is smaller. The TE insertion seems to act as an (unannotated) alternative first exon. In general, several features may be partially covered by a read and a read may extend beyond each of these features. For each pair read-feature we compute the number of bases that are in the read and not the feature (nr) and the number of bases that are in the feature and not in the read (nf). The sum of these two terms nr + nf is the size of the symmetric difference between the two intervals. We assign the read to the feature with the smallest symmetric difference (Figure 1). This situation occurs frequently and assigning a read to a TE only because it covers it yields an overestimation of TE expression (Figure S9 is an example). The impact of each filter is given in Figure 1. After all filters are applied, there are 1 361 (1 301 uniquely mapping (Table S4) in addition to 60 multi-mapping (Table S6)) reads in ovaries and 8 823 (8 551 uniquely (Table S3) and 272 multi-mapped (Table S5)) reads in testes that are assigned to TE copies. This method enables the detection of intergenic TEs, intronic TEs and exonic TEs. Counts are summarised in Table S1.

**Figure 1:**
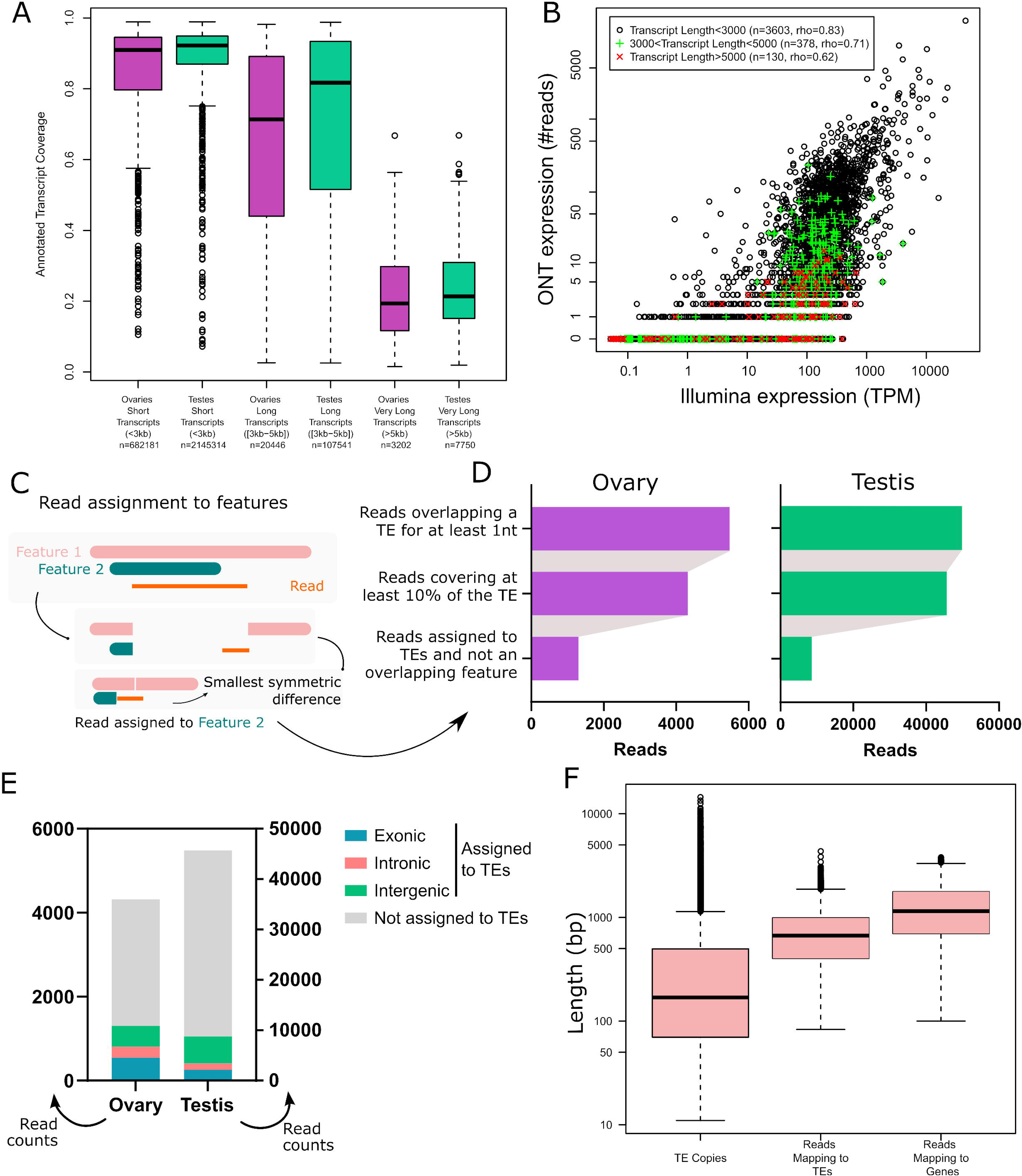
Long-read RNA-seq of *Drosophila melanogaster* ovaries and testes. **A.** Transcript coverage by long-read RNA-seq in ovaries and testes per transcript length (short, long and very long). Very long transcripts (>5Kb) are rare. **B.** Gene expression quantification using Illumina and ONT sequencing in ovaries. Each dot is a gene with a single annotated isoform. Transcripts longer than 5 kb tend to be undersampled using TeloPrime. **C.** Read assignment to features. In the case where a read aligns to a genomic location where two features are annotated, the read is assigned to the feature with the best coverage. Two dimensions are considered. The read should be well covered by the feature, and the feature should be well covered by the read. In practice, we calculate the symmetric difference for each read/feature and select the smallest. In this example, the read is assigned to Feature 2. **D.** Impact of filters on the number of reads assigned to TEs. **E.** Number of reads assigned to TEs separated into three categories (intronic, exonic or intragenic), and reads that overlap TEs but that have not been assigned to TE copies. **F.** TE copy and read length distribution. Reads mapping to TEs encompass most TE copy length but lack transcripts longer than 4.5 Kb, as also observed for reads mapping to genes.

#### Breadth of coverage

To calculate the breadth of coverage of annotated transcripts, we mapped reads to the reference transcriptome and computed for each primary alignment the subject coverage and the query coverage. Scripts used are available on the git repository (sam2coverage_V3.py).

#### Gene ontology

To identify whether ovary and testis had genes associated with their tissue-specific functions, we first selected genes with at least one read aligned in each sample and then we submitted the two gene lists to DAVIDGO separately (Sherman et al., 2022). Due to the high number of biological terms, we selected only the ones with > 100 associated-genes.

#### Subsampling analysis

Subsampling of reads was performed using seqtk_sample (Galaxy version 1.3.2) at the European galaxy server (usegalaxy.eu) with default parameters (RNG seed 4) and the fastq datasets. Subsampled reads were then mapped, filtered and counted using the GitLab/te_long_read pipeline.

#### Splicing

We mapped reads to both the transcriptome and genomic copies of TEs, we selected the ones whose primary mapping was on a TE. We then filtered those exhibiting Ns in the CIGAR strings. Those are the reads aligning to TEs with gaps. We then extracted the dinucleotides flanking the gap on the reference sequence. Scripts used are available on the git repository (SplicingAnalysis.py, splicing_analysis.sh)

### Short-read RNA-seq and analysis

RNA extraction and short-read sequencing were retrieved either from (Fablet et al., 2023), at the NCBI BioProject database PRJNA795668 (SRX13669659 and SRX13669658), or performed here and available at BioProject PRJNA981353 (SRX20759708, SRX20759707). Briefly, RNA was extracted from 70 testes and 30 ovaries from adults aged three to five days, using RNeasy Plus (Qiagen) kit following the manufacturer’s instructions. After DNAse treatment (Ambion), libraries were constructed from mRNA using the Illumina TruSeq RNA Sample Prep Kit following the manufacturer’s recommendations. Quality control was performed using an Agilent Bioanalyzer. Libraries were sequenced on Illumina HiSeq 3000 with paired-end 150 nt reads. Short-read analysis was performed using TEtranscripts (Jin et al., 2015) at the family level, and SQUIRE (Yang et al., 2019) was used for mapping and counting TE copy-specific expression. A detailed protocol on SQUIRE usage in non-model species can be found here https://hackmd.io/@unleash/squireNonModel. Family-level differential expression analysis was performed with TE transcript (Jin et al., 2015). RNA-seq reads were first aligned to the reference genome (GCA_927717585.1) with STAR (Dobin et al., 2013): the genome index was generated with the options --sjdbOverhang 100 and --genomeSAindexNbases 12; next, alignments were performed for each read set with the parameters -sjdbOverhang 100 --winAnchorMultimapNmax 200 and -- outFilterMultimapNmax 100 as indicated by the authors of TE transcript (Jin & Hammell, 2018). TE transcript was ran in two distinct modes, using either multi-mapper reads (--mode multi) or only using single mapper reads (--mode uniq) and the following parameters: --minread 1 -i 10 --padj 0.05 --sortByPos.

## Results and discussion

### Transposable element transcripts are successfully detected with long-read RNA-seq

In order to understand the TE copy transcriptional activity and transcript isoforms in gonads of *D. melanogaster*, we extracted total RNA from ovaries and testes of dmgoth101 adults, a French wild-derived strain previously described (Mohamed et al., 2020). Prior to long-read sequencing, we enriched the total RNA fraction into both capped and polyadenylated mRNAs in order to select mature mRNAs potentially associated with TE activity (see material and methods for the details on the TeloPrime approach). Sequencing yielded between ∼1 million reads for ovaries and ∼3 million reads for testes, ranging from 104 to 12,584 bp (median read length ∼1.4 Kb, Figure S1-2, Table S1). Reads were subsequently mapped to the strain-specific genome assembly (Mohamed et al., 2020) using the LR aligner Minimap2 (version 2.26) (Li, 2018). Most reads mapped to the genome (91.3% for ovaries and 98.8% for testes, Table S1), and the majority of them mapped to a unique location (*i.e.* had no secondary alignment, 98.8% for ovaries and 95.1% for testes), and the vast majority mapped to a unique best location (*i.e.* presence of secondary alignments, one alignment has a score strictly higher than the others, see Methods, 99.9% for ovaries and 97.7% for testes).

In order to validate the long-read RNA-seq approach, we first determined the breadth of coverage of all expressed transcripts and showed that the majority harbour at least one read covering more than 80% of their sequence (70.2% in ovaries and 71.8% in testes). Only a few reads correspond to partially covered transcripts, as most reads cover more than 80% of the annotated transcript sequence (63.4% in ovaries, 77.4% in testes - Figure 1A), although very long transcripts (≥ 5 kb) are poorly covered. The transcriptomes obtained are enriched in typical germline ontology terms, such as “spermatogenesis” for testes, and “oogenesis” for ovaries (Figure S10). Finally, while the first version of the TeloPrime protocol could not be used for quantification (Sessegolo et al., 2019), the quantifications obtained here correlate well with available short-read sequencing (rho=0.78, R=0.44, Figure 1B and S11). We also noticed that the correlation between the two technologies is weaker for very long transcripts.

Although most long reads map to a unique best location on the genome, deciding if a read should be assigned to a TE copy is not straightforward because the read may correspond to a subset or a superset of the annotated TE and it may overlap multiple features (genes, TEs). In this work, we considered the following criteria. First, we only considered reads which cover at least 10% of the annotated TE. Second, when the read overlapped multiple features, we assigned it to the feature for which the coverage was best (see Methods, Figure 1C and Figures S7-8). Our motivation for doing so was to include cases where the read is not TE-only, which is relevant for understanding the broad impact TEs may have in the transcriptome, including old decayed fragmented copies, which may be involved in exonizations, read-through transcripts, upstream promoters, downstream PolyA sites etc (Lanciano & Cristofari, 2020). Restricting the analysis to TE-only reads is also possible using the various metrics we also provide (“Mean % of bases inside TE” and “TE span” on Figure S6 and tables S3 and S4). Overall, after applying these filters, 1 361 reads in ovaries and 8 823 reads in testes were assigned to TE copies (Table S1, Figure 1D). Out of these, 42% are exonic, 20% overlap are intronic and 38% are intergenic in ovaries. In testes, 22% are exonic, 15% intronic and 63% intergenic (Figure 1E).

To check if this long-read dataset is able to recover transcripts encompassing all TE copy lengths present in the genome, we compared the length distribution of all TE insertions with that of all mapped reads (Figure 1F). While genomic TE copies range from a few base pairs to ∼15 Kb, 75% are smaller than 1 Kb. The average length of reads mapping to TEs encompasses the majority of TE copies but does not cover TE transcripts longer than 4.5 Kb. Reads mapping to genes have a similar distribution (Figure 1F). The absence of very long reads (also supported by the cDNA profile, Figure S1) indicates that either very long mRNAs are absent from the sample or the TeloPrime technique is not well tailored for capturing very long transcripts. In order to clarify this point, we compared the quantification obtained by Illumina and ONT TeloPrime for short (<3 Kb), long (3 Kb-5 Kb) and very long transcripts (>5 Kb), and obtained the following Spearman correlations of 0.83 (n=3 603 genes), 0.71 (n=378) and 0.62 (n=130), respectively (Figure 1B for ovaries, Figure S11 for testes). Furthermore, reads covering very long annotated transcripts (>5 kb) tend to be partial (Figure 1A and S12). Therefore, although very long transcripts are rare (<0.1% of reads), the Teloprime protocol could underestimate their presence.

### TE mRNA landscape is sex-specific

One should note that all analysis performed herein take into account all TE annotations in the genome, including old fragmented TE copies. Taking into account all the filtering steps, only 0.3% (8 823/2 925 554) and 0.1% (1 301/1 236 000) of long reads aligned to TE copies in testes and ovaries respectively (Table S1). Given the differences in sequencing depth between both tissues, we have computed the number of reads assigned to TEs based on different sets of subsampled reads, and show that TE reads are more abundant in testes than in ovaries (Figure 2A). We identified 147 TE families supported by filtered long reads (Table S2), of which 78 belong to Long-terminal repeat (LTR) elements (retrotransposons that possess LTR sequences surrounding a retroviral-like machinery). Despite the high number of shared transcribed TE families (101/147), the transcriptional landscape between ovaries and testes is quite different (Figure 2B for the complete dataset and Figure S13 for a subsampled dataset). While LTR elements dominate the transcriptional landscape in both tissues, LINE elements are the second most transcribed TE subclass in testes, while in ovaries, DNA families harbor more read counts (Figure 2B). The transcriptional landscape within TE subclasses between tissues is very similar for LTR retrotransposons, with Gypsy and Pao being the most expressed LTR superfamilies. Jockey retrotransposons dominate the LINE landscape in both tissues, although in ovaries CR1 elements are also observed. The DNA subclass transcriptional landscape is different between testes and ovaries: TcMar-Pogo is the most expressed DNA superfamily in ovaries, while TcMar-Tc1 are abundantly transcribed in testes.

**Figure 2:**
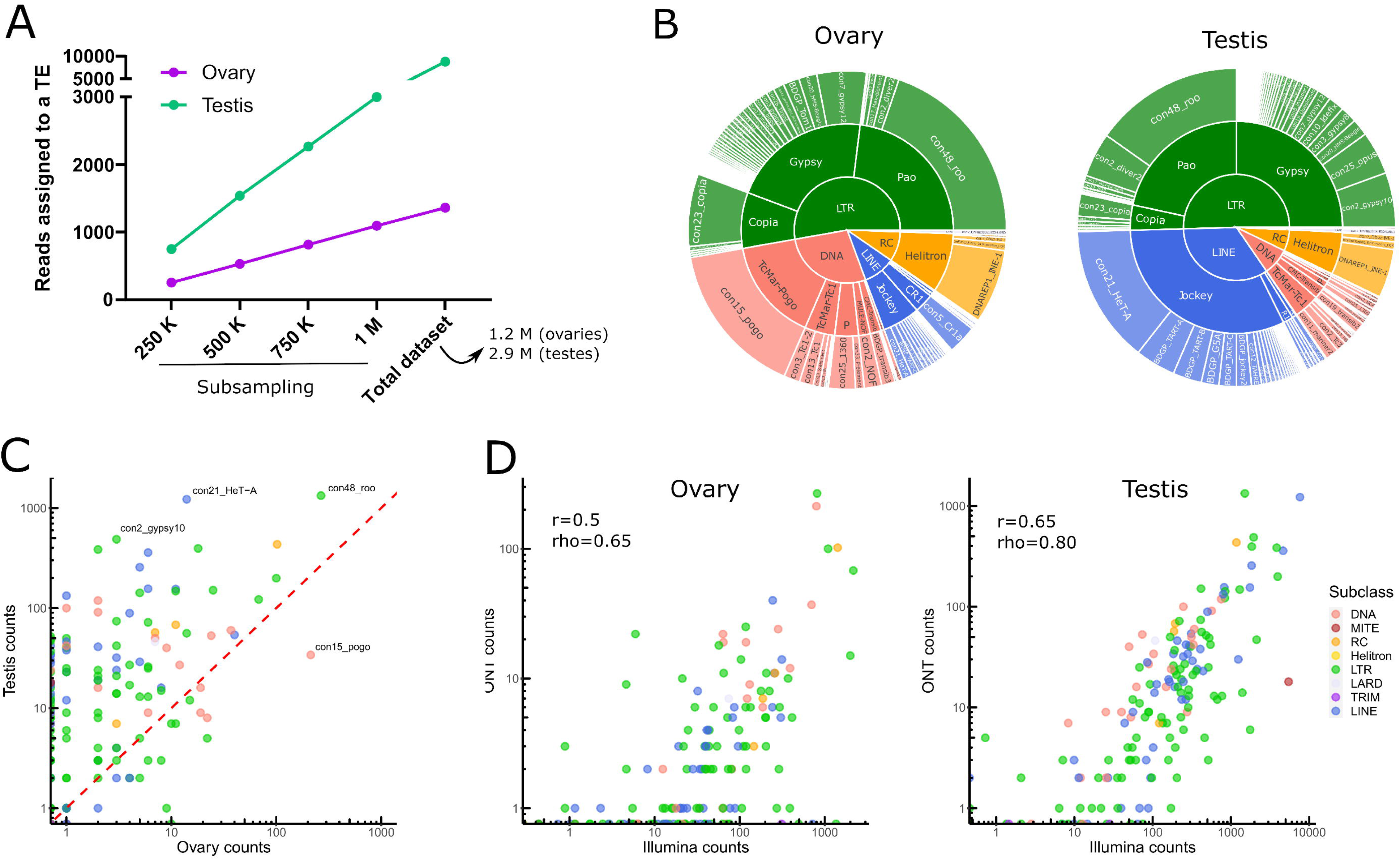
Transposable element transcriptional landscape in ovaries and testes of *Drosophila melanogaster*. **A.** Reads assigned to TEs are more abundant in testis. Subsampling of reads from 250 000 to 1 million reads, along with the complete dataset, show a higher number of reads assigned to TEs in testes than in ovaries. **B.** Global TE transcriptional landscape using ONT long-read sequencing. The outer ring, middle ring and inside circle represent TE family, superfamily and subclass respectively. The area in the circle is proportional to the rate of expression. **C.** Comparison of TE expression ratio between testes and ovaries ONT long-read datasets. **D.** Comparison between Illumina and ONT datasets for estimating the expression levels at the TE family level. Each dot is a TE family. TEtranscript is used for short reads. The correlation between quantifications by both technologies is higher for testes (rho=0.8) than for ovaries, (rho=0.65) where the coverage is weaker. In both cases, it is compatible with what is observed for genes.

Globally, TE families show higher long-read counts in males compared to females (Figure 2C), not only because male samples were more deeply sequenced (2.3 times more), but also because the proportion of reads that map to TEs is higher in males even when subsampling the same number of reads between tissues (Figure S14 for a subsampled dataset). *Con48_roo* (LTR, Pao) and *con21_HeT-A* (LINE, Jockey) are the top two families in male TE long-read counts, with 1331 and 1223 long-reads respectively (Table S2). In females, *con48_roo* (LTR, Pao) and *con15_pogo* (DNA, TcMar-Pogo) are the TE families amounting the most long reads with 266 long reads and 213 (while only 34 in males) respectively (Table S2). There are only four TE families that yielded long-reads in ovaries and not in testes, *con16_blood* (LTR, Gypsy), *BDGP_Helena* (LINE, Jockey), *con9_Bari1* (DNA, TcMar-Tc1), *UnFUnClUnAlig_RIX-comp_TEN* (LINE, I), but most of them harbor only one or two long reads suggesting their expression is low, except *con16_blood* with 10 long-reads. There are 42 families detected only in testes, five DNA elements (*BDGP_transib4* (CMC-Transib), *con3_looper1* (con3_looper1), and three TcMar-Tc1 (*BDGP_Bari2, con9_UnFUnCl001_DTX-incomp* and *con48_FB*), LINE elements (11 Jockey, two R1, and one CR1, see Table S2 for details), 21 LTR families (two Copia, Gypsy, and three Pao), and one MITE family, ranging from one to 52 long reads per TE family. Finally, 17 TE families show no long-read mapping in either tissue. Collectively, long-read sequencing can discriminate between ovaries and testes TE transcriptional landscapes, however a robust analysis supporting these differences would require replicating the results.

Short-read RNA sequencing of ovaries and testes, followed by estimation of TE family expression with TEtranscripts (Jin et al., 2015) - TE expression estimation *per* TE family, see material and methods for more information) recapitulates the long-read RNA sequencing profiles (Figure 2D and Figure S15). Although TE transcripts are overall poorly expressed, the estimation of their expression level is reproducible across technologies. The correlation is higher for testes (r=0.65, rho=0.8) than for ovaries (r=0.5, rho=0.65), where the coverage is weaker. Indeed, as previously stated, the total contribution of TEs to the transcriptome is weaker for ovaries and the sequencing is shallower.

### Long-read sequencing successfully retrieves copy-specific transcripts

The main objective of using long-read sequencing after the TeloPrime full-length cDNA enrichment protocol is to recover copy-specific mature TE transcripts and potential isoforms. There are 1 301 long reads mapping uniquely to a TE copy in ovaries, while 52 map to multiple copies within the same family and eight reads are not assigned to a specific TE family. In testis, 8 551 reads are assigned to specific TE copies, 200 to TE families and 72 are assigned at the superfamily or subclass level. The overall percentage of reads unable to be assigned to a particular copy is therefore quite small (4.4% and 3% for ovaries and testes respectively). The only family harboring only multimapped reads is *con10-accord* with one single long-read in ovaries that matches two different copies.

In ovaries, out of 105 TE families detected (at least one read), 13 families harbor only one multi-mapped read, and six families have 2 to 21 multimapped reads (Figure 3A). While only ∼4% of *con15_pogo* reads are multimapped in ovaries (8 out of 213 reads), *con23_copia* harbor a higher percentage of multimapped reads, 21% of 100 reads for *con23_copia.* In testis, 143 TE families are expressed (at least one read), and 43 have multi-mapped reads (Figure 3A). As observed in the ovary dataset, the number of multimapped reads is low for most families, with only six families harboring more than 10 multimapped reads. While *con23_copia* harbors the most multimapped reads in testis (46 out of 199 long-reads), *con3_looper1* and *con20_Burdock* show a higher multi-mapped read ratio with ∼58% of reads multimapped out of 24 and 19 reads respectively. In total, 50 TE families have both uniquely and multi-mapping reads in ovaries and testis (Figure 3A).

**Figure 3:**
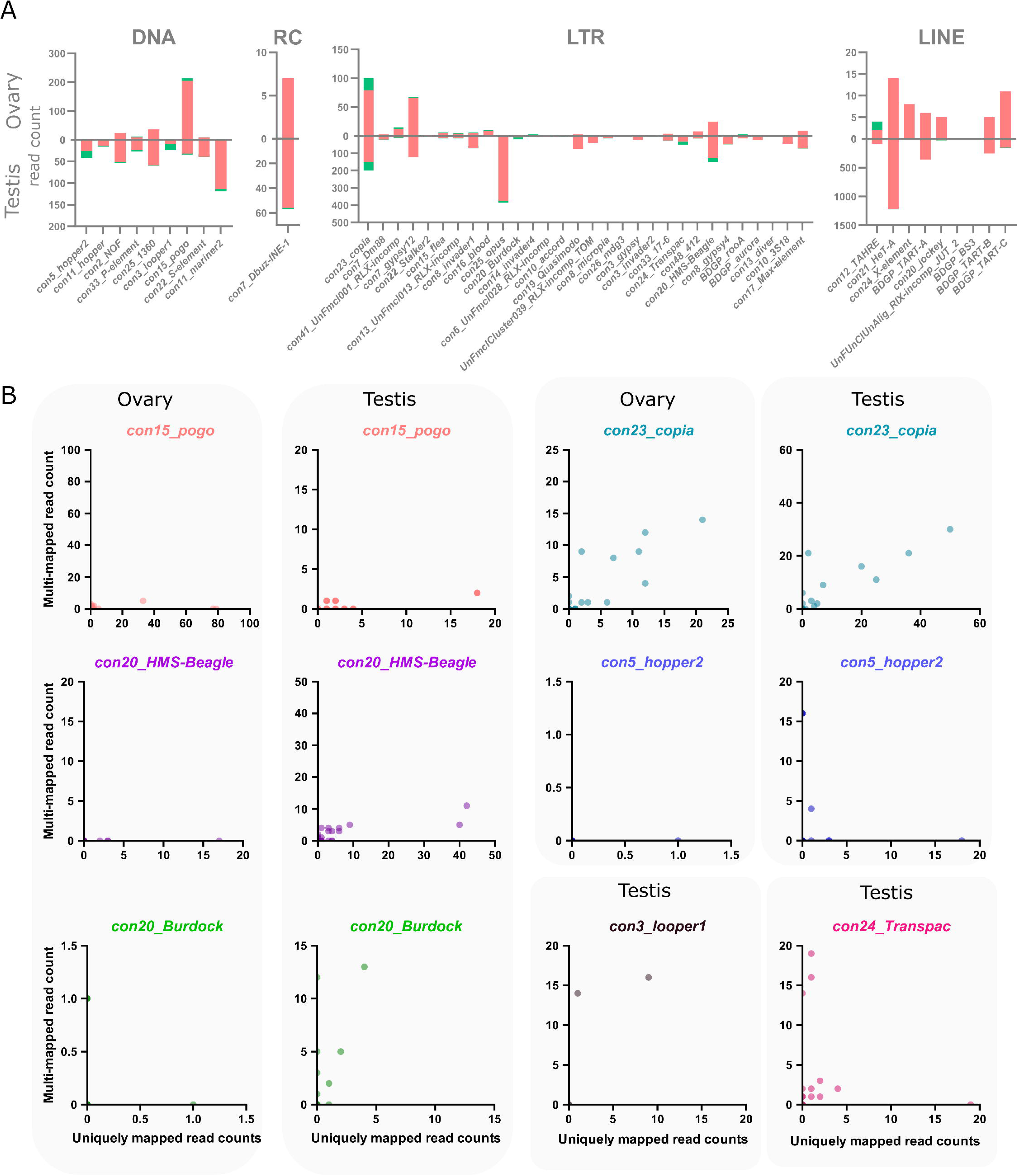
Multi-mapping and uniquely mapping ONT reads. **A.** Distribution of uniquely and multimapped reads across TE families in ovaries and testes (only TE families harboring at least one multimapped read are shown), see Figure S16 for a figure including *con48_roo*. **B.** Association between multi-mapped and uniquely mapped reads at the copy-level for the TE families showing a high number of multi-mapped reads. Each dot represents a unique genomic copy, for *con3_looper1* and *con24_Transpac*, only testis samples are shown as no copies were expressed in ovaries.

We uncovered 443 and 1 165 TE copies harbouring at least one long-read unambiguously mapping in ovaries and testes respectively. When taking into account multi-mapped reads, an additional 55 and 89 TE copies are potentially expressed in each tissue (Table S2). However, it is important to note that the number of assigned multi-mapped reads to each copy is quite small, as seen at the family level. For instance, in ovaries, 46 of these potentially expressed copies only harbor one multi-mapped read, 16 copies harbor two, and a single *pogo* copy harbors three multi-mapping reads. As a comparison, the two most expressed copies in ovaries are two *con15_pogo* copies, con15_pogo$3L_RaGOO$9733927$9735150 and con15_pogo$X_RaGOO$21863530$21864881, with 79 and 77 uniquely mapping reads, and no multi-mapping read (Table S4). In testis, out of 89 copies without uniquely mapping reads, 65 have only a single multi-mapped read, and only seven copies show more than 10 multi-mapped reads (Table S3). As a comparison, the top expressed copy in testis is a con2_*gypsy10* (con2_gypsy10$3R_RaGOO$760951$766941) with 472 uniquely mapping reads and no multimapping ones. Finally, among TE families showing the highest number of multimapping counts, *con23_copia* tops with 21 multimapped reads on ovaries and 46 in testis, nevertheless, with the exception of two copies harboring one or two multimapping reads, all other detected *con23_copia* insertions show uniquely mapping reads (Figure 3B). There are however a few TE families where the identification of single-copy transcripts is hazardous. For instance, *con5_hopper2* has one copy clearly producing transcripts with 18 uniquely mapping reads and no multimapping ones, however, there are three other copies that share 16 multimapped reads and no uniquely ones (Table S3, Figure 3B). Therefore, despite a few exceptions, long-read sequencing can identify single-copy TE transcripts.

Within a TE family, the contribution of each TE copy to the family transcriptional activity is variable. In general, only a few insertions produce transcripts, even if taking into account multi-mapped reads (Figure 4A for uniquely mapping reads and Figure S17 for all reads). However, *con24_Transpac* (LTR, Gypsy) copies are nearly all expressed in testes (8 expressed copies and four potentially expressed copies out of 16), while in ovaries, *con15_pogo* (DNA, TC-mar-Pogo) harbors 13 copies producing transcripts, and six potentially expressed copies out of 57. *DNAREP1_INE-1* (RC) is the most abundant TE family in the *D. melanogaster* genome and is also the family harbouring the most transcribed copies in both ovaries and testes (51 and 118 respectively out of 1 772 copies). The *con48_roo* also show many expressed insertions with 69 in ovaries and 112 in testis out of 475 copies. Finally, in ovaries, out of the 443 insertions with at least one mapped read, 25 had more than 10 mapped reads. In testes, out of the 1 165 insertions with at least one uniquely mapped read, 160 had in fact more than 10 mapped reads.

**Figure 4:**
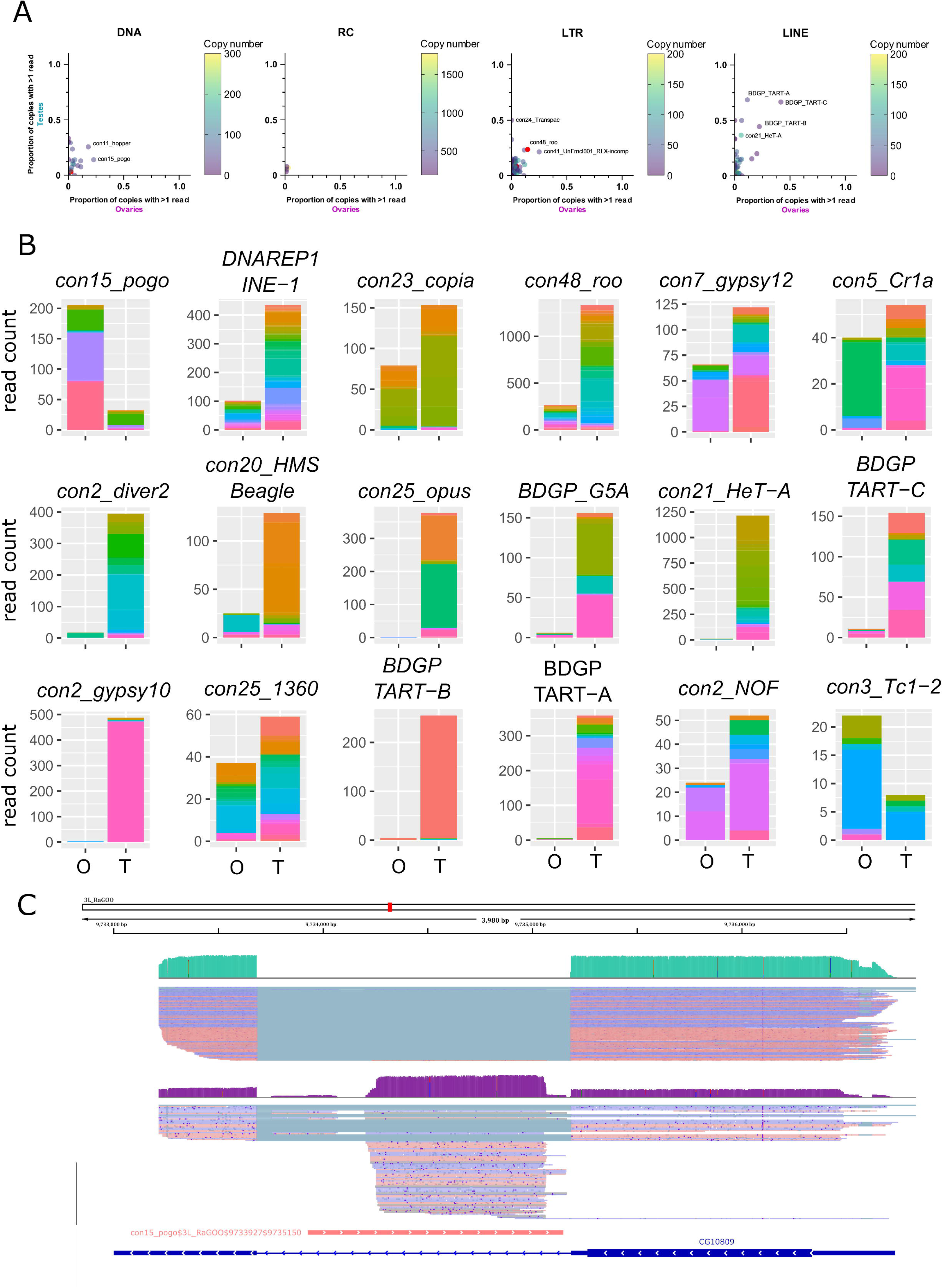
Transcription of transposable element copies. **A.** Frequency of transcribed copies (read > 1) within TE subfamilies in ovaries and testes, along with genomic copy number (color bar, 1 to 200 (LINE/LTR) or 300 (DNA) copies). All TE families harbouring more than 200 (LINE/LTR) or 300 (DNA) copies are depicted in pink. For DNA elements, *con25_1360* has 875 insertions. For LINE families, *con5_Cr1a* has 671 copies. The LTR families, *con10_idefix* (251), *UnFmclCluster039_RLX-incomp_COR* (278), *con19_Quasimodo* (253), *con2_diver2* (205), *con48_roo* (475), *con7_gypsy12* (242) and *con3_gypsy8* (315) are also depicted in pink. Most TE subfamilies have only a couple of copies producing transcripts, while the majority of HETA copies are expressed in testes for instance (middle panel). **B.** Distribution of read counts per copy for the 10 most expressed families in ovaries and testes (16 TE families total), showing the overall expression of specific copies within a TE family (Table S3 and S4). Copies are represented by different colors within the stacked bar graph. Uniquely mapped reads are used. O: ovaries, T: testes **C.** IGV screenshot of a *pogo* copy (con15_pogo$3L_RaGOO$9733927$9735150, in pink). In green, testis coverage and below mapped reads, in purple the same information for ovaries. Dmgoth101 repeat and gene tracks are also shown and more information on the annotation can be seen in the material and methods section.

While many TE copies within a family produce transcripts, there are significant differences in copy expression rate (Figure 4B for the 10 most expressed TE families in ovaries and testes, and Figure S18 for a subsampled dataset). For instance, *con25_1360* (DNA, P) has many transcriptionally active copies, and a similar activity landscape between ovaries and testes. While many copies produce transcripts within the *con5_Cr1a* family, the transcriptionally active copies differ between the tissues studied. *Con2_gypsy10* (LTR - Gypsy) harbors a highly active copy with 472 uniquely mapping reads out of 488 total counts in testes, while only three reads are detected in ovaries (Figure 4B).

In ovaries, where *con15_pogo* has one of the highest number of long reads (213), an insertion of 1 222 bp in the 3L chromosome (con15_pogo$3L_RaGOO$9733927$9735150) accumulates nearly 37% of the family total read count (Figure 4B and C). This specific *pogo* insertion is located in the intron of the CG10809 gene. The same pattern is observed for the second most expressed *pogo* insertion (con15_pogo$X_RaGOO$21863530$21864881), also located in the intron of a gene (CG12061), expressed in testes and not in ovaries (Figure S19). CG12061 is a potential calcium exchange transmembrane protein and has been previously shown to be highly expressed in the male germline (Li et al., 2022). Indeed, using long-read sequencing, CG12061 is highly expressed in testes compared to ovaries, and curiously, the intronic *pogo* insertion is only expressed when the gene is silent (in ovaries). The other expressed insertions of *pogo* are located in intergenic regions (Figure S20).

Finally, using short-read sequencing and a tool developed to estimate single-copy expression (Squire (Yang et al., 2019)), we compared the overall TE copy transcriptional landscape between short and long reads (Figure S21). There was a poor correlation with the ONT estimations (rho=0.23, r=0.28 for ovaries and rho=0.36, r=0.32 for testes). At the family level, the quantifications obtained by Squire were comparable to the ones obtained with long-reads (rho=0.59, r=0.71 for ovaries and rho=0.77, r=0.66 for testes, Figure S21). Examination of instances where the two techniques differed most, shows that Squire tends to overestimate the expression of TE insertions completely included in genes (Figure S9). Indeed, while long-reads can easily be assigned to the correct feature because they map from the start to the end of the feature, many of the short-reads originating from the gene also map within the boundaries of the TE. Methods based on short reads could clearly be improved, based on the study on such instances where there is a discordance.

### Transcripts with TE sequences may extend beyond annotated TE boundaries

The previous analysis focused on reads that cover 10% of the TE sequences and have been assigned to TE copies by the “feature assignment” filter (see Methods). This analysis includes reads that potentially align beyond the TE boundaries. In practice, this is often the case, as ⅔ of transcriptional units associated to TEs are covered by reads which, on average, have more than 10% of their sequence located outside the TE (Figure S6). If we restrict our analysis to TE copies covered by reads with more than 90% of their sequence inside the TE, we obtain, in ovaries, 128 expressed TE copies supported by 491 uniquely mapping reads, and in testes, 323 expressed TE copies, supported by 2 969 uniquely mapping reads. The TE transcriptional landscape (Figure S22) is similar to the one obtained with the default filters (Figure 2B). The main difference is the absence of *con48_roo and DNAREP1_INE-1.* Indeed, many cases of expressed *con48_roo* insertions correspond to TE-gene chimeras (Figure S23, Figure S24). The reads overlap both the TE and the gene but our algorithm assigns them to the TE because they overlap more the TE than the gene (smallest symmetric difference, see Figure 1C and methods). In some other cases however, there is no annotated gene and the reads still extend beyond the TE boundaries. This is the case of *con2_diver2* (Figure S25). In contrast, *con15_pogo, con23_copia, con2_gypsy2* are among the most expressed families even when using these more stringent filters, suggesting these families are transcriptionally active.

### Transcripts from full-length transposable element copies are rarely detected

Many insertions produce transcripts that are shorter than the annotated TE and are likely unable to participate in TE transpositional activity. Furthermore, even in the case where the transcript fully covers the insertion, the copy itself might have accumulated mutations, insertions and deletions making it unable to transpose. To assess this, we computed the query coverage of the reads with regards to the insertion they correspond to. We find that one-third of the insertions have at least 80% query coverage (Figure 5A). However, out of these insertions, only a few of them are close in length to a functional full-length sequence. In order to search for potentially functional, expressed copies, we filtered for copies with at least five long-reads detected, and covering at least 80% of their consensus sequences. In ovaries these filters correspond to eight insertions: one con15_*pogo*, six *con23_copia* insertions and one *con17_Max*, and in testes, there are eight potentially functional insertions, five *Copia* insertions, one *BDGP_Bari2*, and one *con6_1731*. While all *con23_copia* insertions expressed with at least five reads are full-length, other TE families show mostly internally deleted expressed copies (Figure 5B). Indeed, a closer analysis of *con15_pogo*, the most expressed TE subfamily in ovaries, shows only one full-length copy expressed (con15_pogo$2L_RaGOO$2955877$2958005), but at low levels (five reads in ovaries, and two in testes). Instead, the other three expressed *con15_pogo* copies with at least five reads in ovaries (79, 77 and 33), are internally deleted (Figure 5B). Hence, ONT long-read sequencing detects only a small number of expressed full-length copies. As stated before, very few cDNA molecules longer than ∼4 Kb have been sequenced (Figure S1), suggesting either that such longer transcripts are rare, and/or that the method used here for cDNA amplification induces a bias towards smaller sequenced fragments. Expression of longer TE copies might therefore be underestimated.

**Figure 5.**
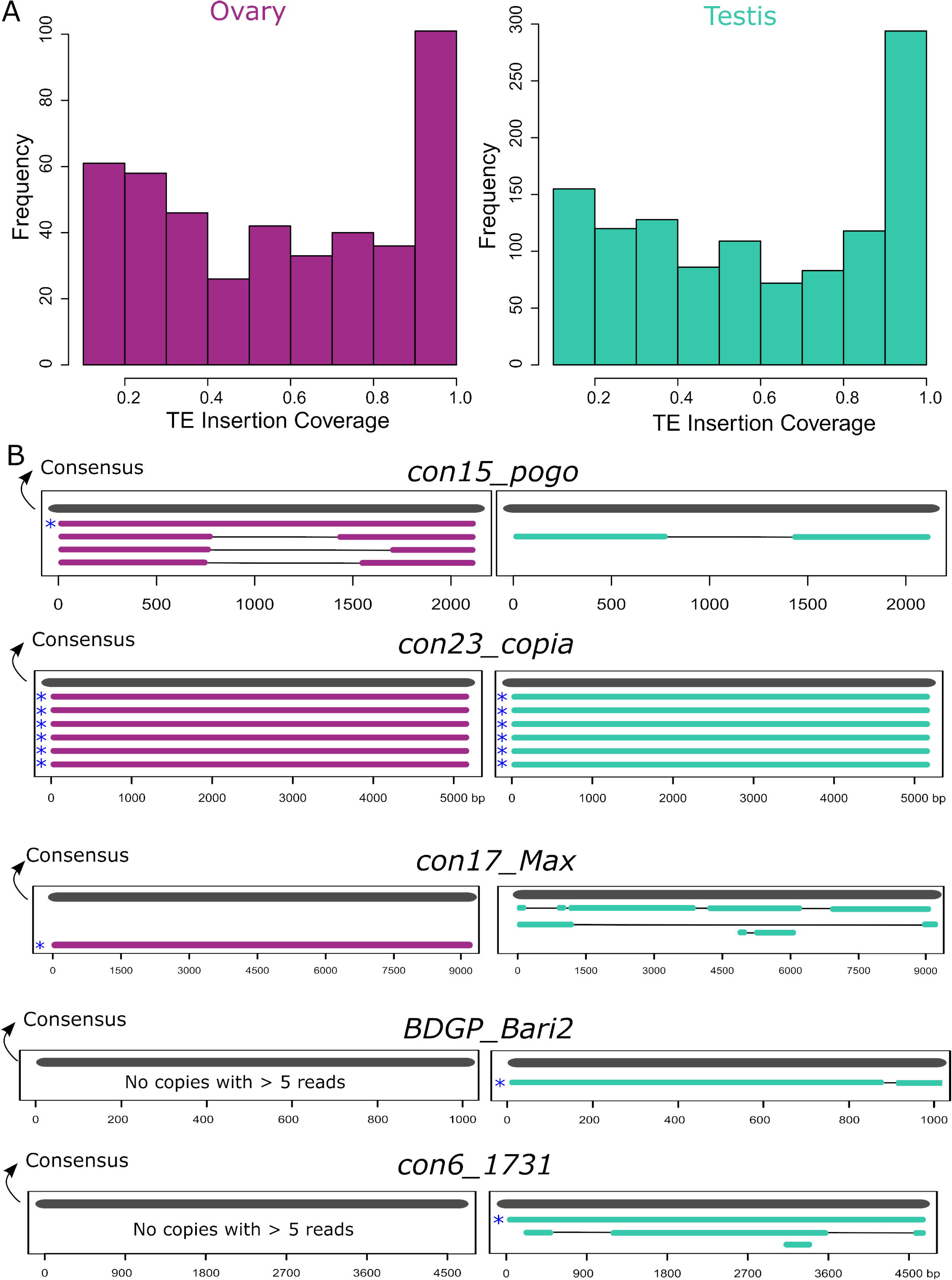
TE transcripts stem mostly from deleted or truncated copies. **A.** Coverage of ONT reads on TE insertions. One-third of copies are covered for at least 80% of their length. **B.** Alignment of copies belonging to the five TE families where at least one full-length expressed copy (80% of consensus) was observed with more than five long-reads. All copies represented have at least five long-reads. Consensus sequences are represented in grey and copies are either purple for ovaries or green for testes. Asterisks depict the full-length copies.

### Long-read sequencing may unvail novel spliced TE isoforms

A closer analysis of the reads stemming from the detected full-length copies shows that many of them do not cover the copies completely (Figure 6). For instance, six *con23_copia* copies are at least ∼80% of the consensus sequences and have at least five long-reads detected (Figure 5B), however, although the reads map from the 5’ end to the 3’ end of the copy, they map with a gap (Figure 6 and Figures S26-S28). *con15_pogo, con17_Max-element, con6_1731* also show such gapped alignments. Inspection of these gaps reveals that they are flanked by GT-AG consensus, suggesting that those transcripts are spliced. In contrast, *BDGP_Bari2* shows five reads that correspond to the full-length copy and three that extend beyond the TE boundaries. Out of these three reads, one aligns with a gap, flanked by GT-AG, the gap itself overlaps the TE boundaries. One should note that the consensus sequence of *BDGP_Bari2* is small (1kb), while the other elements are much longer. Collectively, long-read sequencing shows that despite the presence of potentially functional, full-length copies in the *D. melanogaster* genome, only a few of these are detected as expressed in testes and ovaries, and the reads that are indeed recovered seem to be spliced.

**Figure 6.**
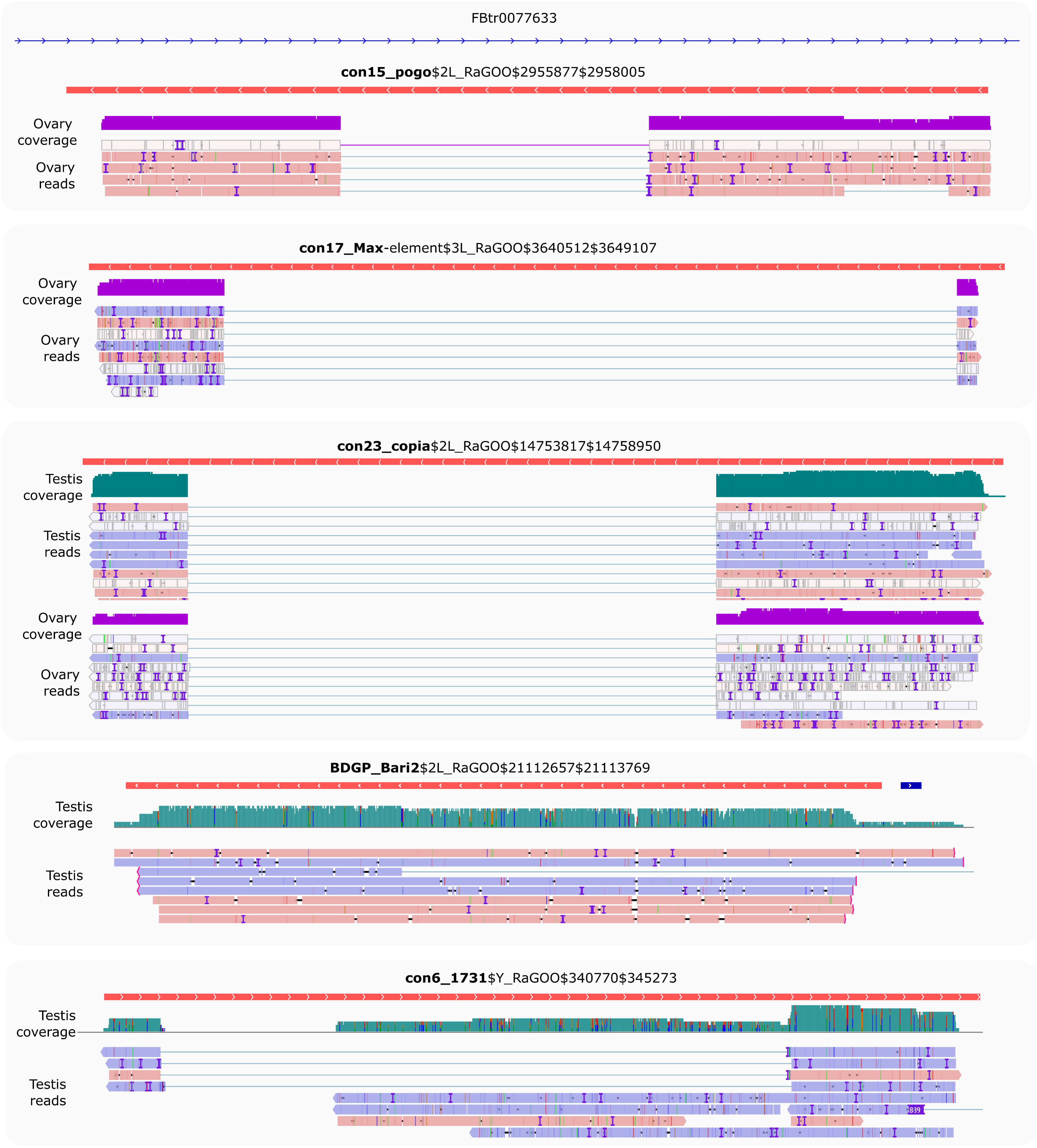
Full-length copies produce spliced transcripts. IGV screenshot of uniquely mapping reads against putative full-length copies (copies > 80% of the consensus sequence length) harboring at least five reads (see Figure 5B). Only *con23_copia* has multi-mapped reads that can be appreciated in Figure S26. Dmgoth101 repeat and gene tracks are also shown and more information on the annotation can be seen in the material and methods section. Ovary and/or testis coverage and reads are shown below the TE copies.

While most TEs do not harbor introns, there are a couple of exceptions previously described in *D. melanogaster.* Indeed, P elements are known to be regulated tissue-specifically by alternative splicing mechanisms, involving piRNA targeting (Laski et al., 1986; Teixeira et al., 2017). *Gypsy* copies are able to produce ENV proteins through mRNA alternative splicing (Pélisson et al., 1994; Teixeira et al., 2017). As with P elements, *gypsy* splicing is also thought to be regulated by piRNAs. Finally, *Copia* elements produce two isoforms, a 5 Kb and a 2.1 Kb (which is a spliced product of the 5 Kb mRNA) (Miller et al., 1989; Yoshioka et al., 1990). The 2.1 Kb encodes the GAG protein and is produced at higher levels than the other proteins (Brierley & Flavell, 1990). While the shorter transcript can be processed by *Copia* reverse transcriptase, the 5 Kb full-length isoform is clearly preferred (Yoshioka et al., 1990). Most of these discoveries were obtained through RT-PCR sequencing of amplicons, or recently, through short-read mapping. Nevertheless, systematic analysis of TE alternative splicing in *D. melanogaster* is lacking, due to the difficulty of detecting such isoforms from short-read data. Here we used long-read sequencing to mine for such splicing isoforms. We searched for reads harboring a gap compared to the reference sequence (presence of N’s in the CIGAR string). In order to ensure that those gaps corresponded to introns, we searched for flanking GT-AG splice sites (see methods, and Figures S29-S32). In ovaries, out of 25 insertions supported by at least five reads, 17 exhibited at least one gapped read (Figure 7, Table S8). Out of these 17 cases, 13 corresponded to GT-AG consensus. The four remaining cases were 1 CT-AC, 1 CT-TA, 1 CT-TG, 1 TA-GT. In testes, out of 201 insertions supported by at least five reads, 112 exhibited at least one gapped read, 59 with a GT-AG consensus, 43 with a CT-AC consensus (Figure 7, Table S7). Out of the 10 others, seven exhibited only one or two gapped reads, and the three remaining were GC-AG (con14_Rt1a$2R_RaGOO$3576319$3576878), CT-AT (con8_UnFmcl025_RLX-comp$2R_RaGOO$2841061$2845392) and AC-CG (BDGP_G5A$2R_RaGOO$4442347$4444567). Those could correspond to non-canonical splicing, to a heterozygous deletion, or to the expression of a deleted copy located in a non-assembled part of the genome.

**Figure 7.**
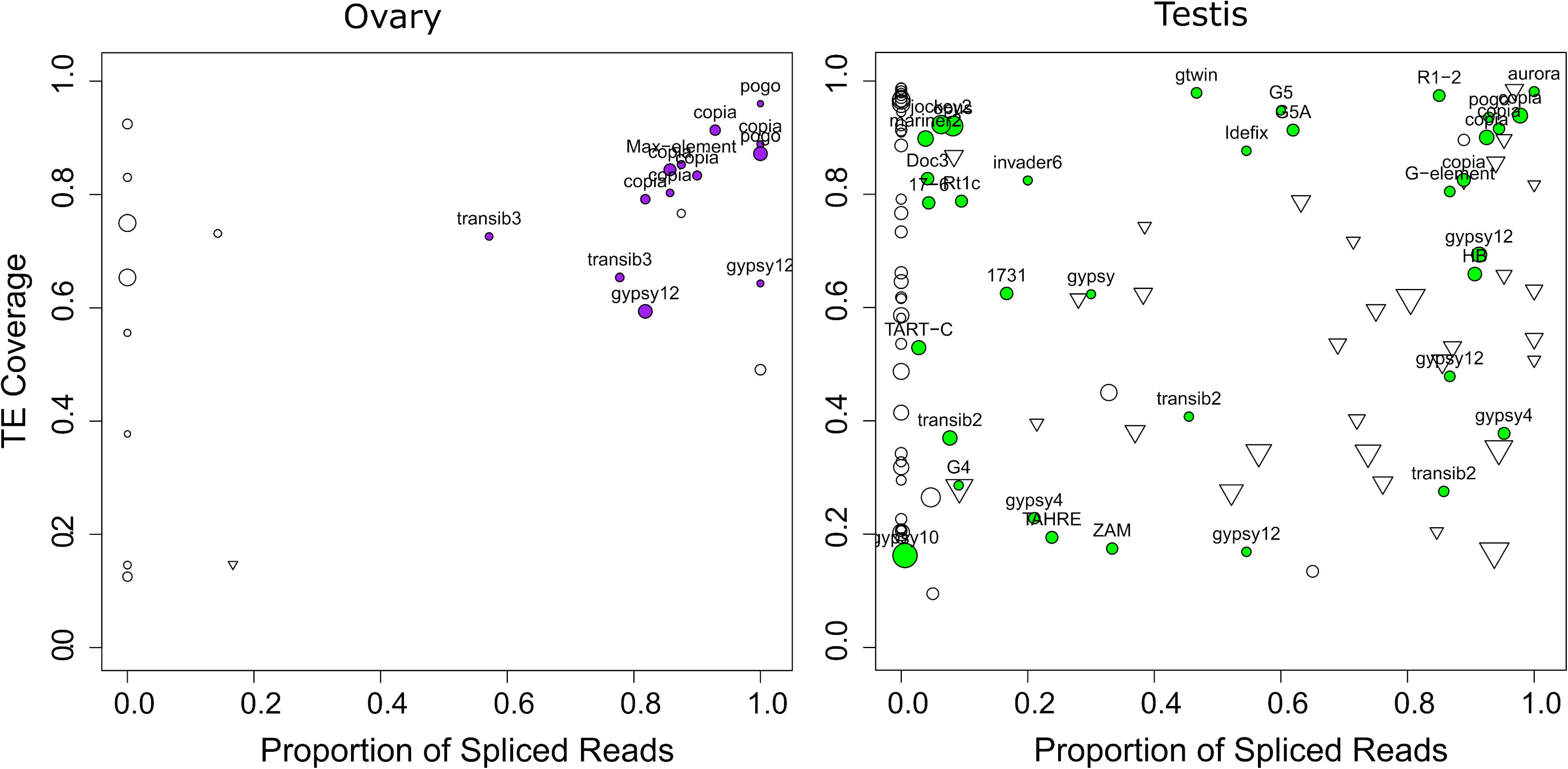
TE spliced transcripts are frequent. Left depicts ovaries, right depicts testes. Each circle depicts a TE insertion supported by at least 5 reads in ovaries (resp. 10 reads in testes) and their size is proportional to the expression level of the insertion. The TE family name is written for gaps with a GT-AG consensus. The X-axis represents the proportion of reads that align with a gap (presence of N’s in the CIGAR string), while the Y-axis represents the proportion of the insertion covered by reads. Inverted triangles correspond to gaps with a CT-AC consensus. Unfilled circles correspond either to TE insertions with no gaps, or to TE insertions with gaps that do not exhibit either GT-AG or CT-AC sites.

The fact that we identify gaps with a CT-AC consensus suggests that the pre-mRNA that is spliced is transcribed antisense with respect to the TE. Our long reads are not stranded, but we could verify using our stranded short-read data, that the transcription was indeed antisense (Figure S33). This verification was possible for cases where there were enough uniquely-mapped short reads. This was however rarely the case as the TE copies containing CT-AC gaps had recently diverged. Out of the 43 CT-AC instances, the most represented families were *con21_HeT-A* (15), *BDGP_TART-A* (6), *BDGP_TART-C* (5). These TEs are involved in telomeric DNA maintenance. Antisense transcription has already been reported for these elements (Casacuberta & Pardue, 2005), which is coherent with our hypothesis that spliced antisense transcripts are also captured. Overall, we find that the majority of gaps are flanked by GT-AG consensus (or CT-AC), and we conclude that they correspond to spliced introns. These introns are however not systematically spliced, because in many cases the proportion of spliced reads is between 0 (never spliced) and 1 (always spliced).

While the proportion of spliced transcripts stemming from a TE copy can vary, there are a couple of copies that only produce spliced transcripts, as con15_pogo$2R_RaGOO$7201268$7202755 for instance. *con15_pogo* is the most expressed TE family in ovaries, with 13 out of 57 copies producing capped poly-A transcripts corresponding to 213 long reads, while only 8 expressed copies with a total of 34 long-reads are observed in testes, despite the higher coverage. While we previously noted that only one full-length copy is transcribed in ovaries (and in testes albeit with a lower number of reads), there are many truncated or deleted copies that are transcribed (Figure 5B). con15_pogo$2R_RaGOO$7201268$7202755 is one of the internally deleted copies, and it produces a spliced transcript present in both testis and ovaries (Figure S20). The splicing of this short intron (55 nt) has been previously reported (Tudor et al., 1992) and enables the splicing of the two ORFs of *pogo* into a single continuous ORF. This particular copy (con15_pogo$2R_RaGOO$7201268$7202755) is however non-functional since it contains a large genomic deletion located in the ORF near the intron. con15_pogo$X_RaGOO$21863530$21864881 (Figure S19) also contains a large genomic deletion, encompassing the intron, explaining why there are no spliced transcripts for this copy.

Despite the presence of full-length *con23_copia* insertions in the genome, only spliced transcripts were uncovered in the long-read sequencing (Figure 6). In contrast, with Illumina short reads, we see both spliced and unspliced transcripts (Figure 8). A similar pattern occurs with con17_Max-element$3L_RaGOO$3640512$3649107 (Figure S34), but not with con15_pogo$2L_RaGOO$2955877$2958005 or con6_1731$Y_RaGOO$340770$345273 (Figures S35-36). The full-length *con23_copia* transcripts are 5 Kb, and are less abundant than the spliced transcripts (∼10 times less). The lack of such a full-length transcript in the long-read sequencing data might be explained by the lower expression level and the length of the transcript. One can not discard the possibility that deeper long-read coverage might uncover full-length, unspliced, *con23_copia* transcripts. It is important to stress that by using only short-reads it is nearly impossible to determine which *con23_copia* sequence is being expressed as the vast majority of short reads map to multiple locations with the exact same alignment score. With short-reads, at least one full-length *con23_copia* insertion is expressed but its specific location remains unknown. Furthermore, if we restrict the analysis to primary alignments (i.e. a randomly chosen alignment in the case of multiple mapping), then the coverage of the intronic sequence decreases and it is no longer clear if the insertion produces both spliced and unspliced transcripts (Figure S27). Overall, for *con23_copia*, the long-reads enable the identification of which insertion is being transcribed, and the short-reads enable the detection of the presence of the two splice variants. Some multi-mapping long reads could support the presence of the unspliced transcript because they partially map to *con23_copia* intron, but we cannot know from which insertion they were transcribed (Figure S28). Finally, spliced transcripts are unable to produce the complete transposition machinery as they lack the reverse transcriptase enzyme and are only able to produce the gag protein.

**Figure 8.**
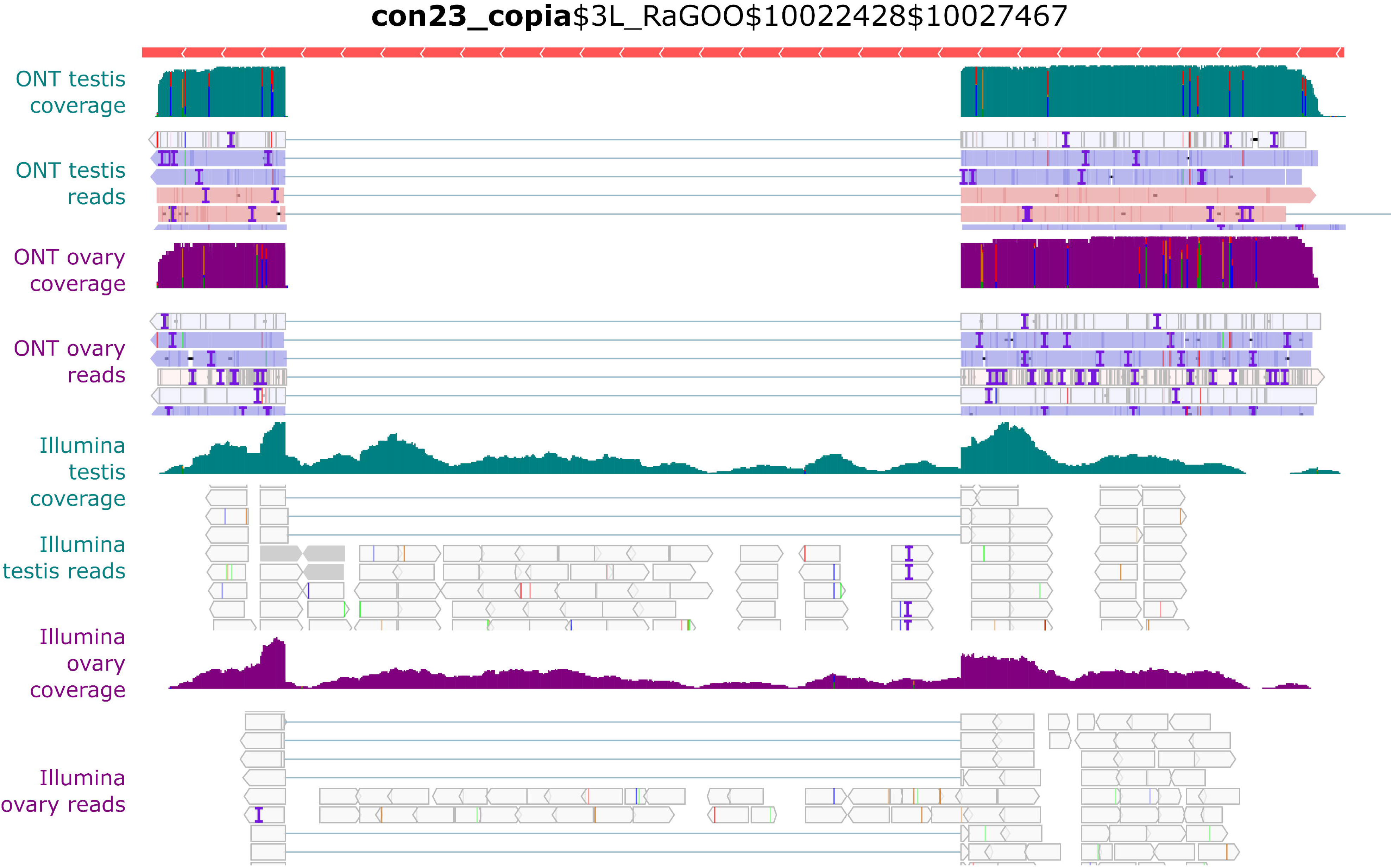
Example of *con23_copia* splicing. IGV screenshot shows spliced transcripts using long and short-read datasets. In green, testis coverage and an excerpt of mapped reads, in purple the same information for ovaries. In the excerpt of mapped reads, white rectangles correspond to multi-mapping reads. Dmgoth101 repeat and gene tracks are also shown and more information on the annotation can be seen in the material and methods section.

## Conclusion

Long-read sequencing largely facilitates the study of repeat transcription. Here we demonstrated the feasibility of assigning long reads to specific TE copies. In addition, quantification of TE expression with long-read sequencing is similar to short-read analysis, suggesting not only one could recover copy-specific information but also perform quantitative and differential expression analysis.

The genome of *D. melanogaster* contains many functional full-length copies but only a couple of such copies produce full-length transcripts in gonads. Given TEs are tightly controlled in the germline, one can wonder how many full-length copies might be expressed in somatic tissues. It is also important to stress that, to our knowledge, this is the first comparison of the expression of TEs between testes and ovaries, and we uncover a different TE transcriptional landscape regarding TE subclasses, using both short-reads and long-reads. Furthermore, in many instances, we see that TE transcripts are spliced, independently of their structure or class. While some of these introns had been reported in the literature 30 years ago, the relevance and prevalence of these spliced transcripts have not always been investigated. Long-read sequencing could facilitate the exhaustive inventory of all spliceforms, in particular for recent TEs, where short reads are harder to use due to multiple mapping. While our results suggest that TE splicing could be prevalent, additional studies with biological replicates, high sequencing coverage and mechanistic insights into the splicing machinery will be needed to confirm our observations. A difficulty that remains when assessing if the intron of a particular TE insertion has really been spliced is the possibility that there exists a retrotransposed copy of a spliced version of this TE elsewhere in a non-assembled part of the genome. Here, taking advantage of the availability of raw genomic Nanopore reads for the same dataset (ERR4351625), we could verify that this was not the case for *con23_copia*, the youngest expressed element in our dataset. In practice, we mapped the genomic reads to both *con23_copia* and a spliced version of *con23_copia* and found no genomic read mapping to the spliced version.

Finally, it is important to note that we did not recover TE transcripts longer than ∼4.5 Kb. While the detection of rare transcripts might indeed pose a problem to most sequencing chemistries, it would be important to verify if long transcripts necessitate different RNA extraction methods for ONT sequencing. For instance, the distribution of cDNA used here for ONT library construction reflects the distribution of reads, with a low number of cDNAs longer than 3.5 Kb (Figure S1).

## Acknowledgements

This work was performed using the computing facilities of the LBBE/PRABI. We thank Josefa Gonzalez laboratory for discussing our preprint in their journal club and suggesting modifications to the manuscript.

## Data, scripts, code, and supplementary information availability

Data are available online at the BioProject PRJNA956863 (ONT long-reads), PRJNA981353 (SRX20759708, SRX20759707, testes short-reads), PRJNA795668 (SRX13669659 and SRX13669658, ovaries short-reads). Scripts are available at https://gitlab.inria.fr/erable/te_long_read. Processed data (.bam files) are available at https://zenodo.org/records/10277511.

