## Supplementary Figures for "Identification and quantification of transposable element transcripts using Long-Read RNA-seq in *Drosophila* germline tissues"

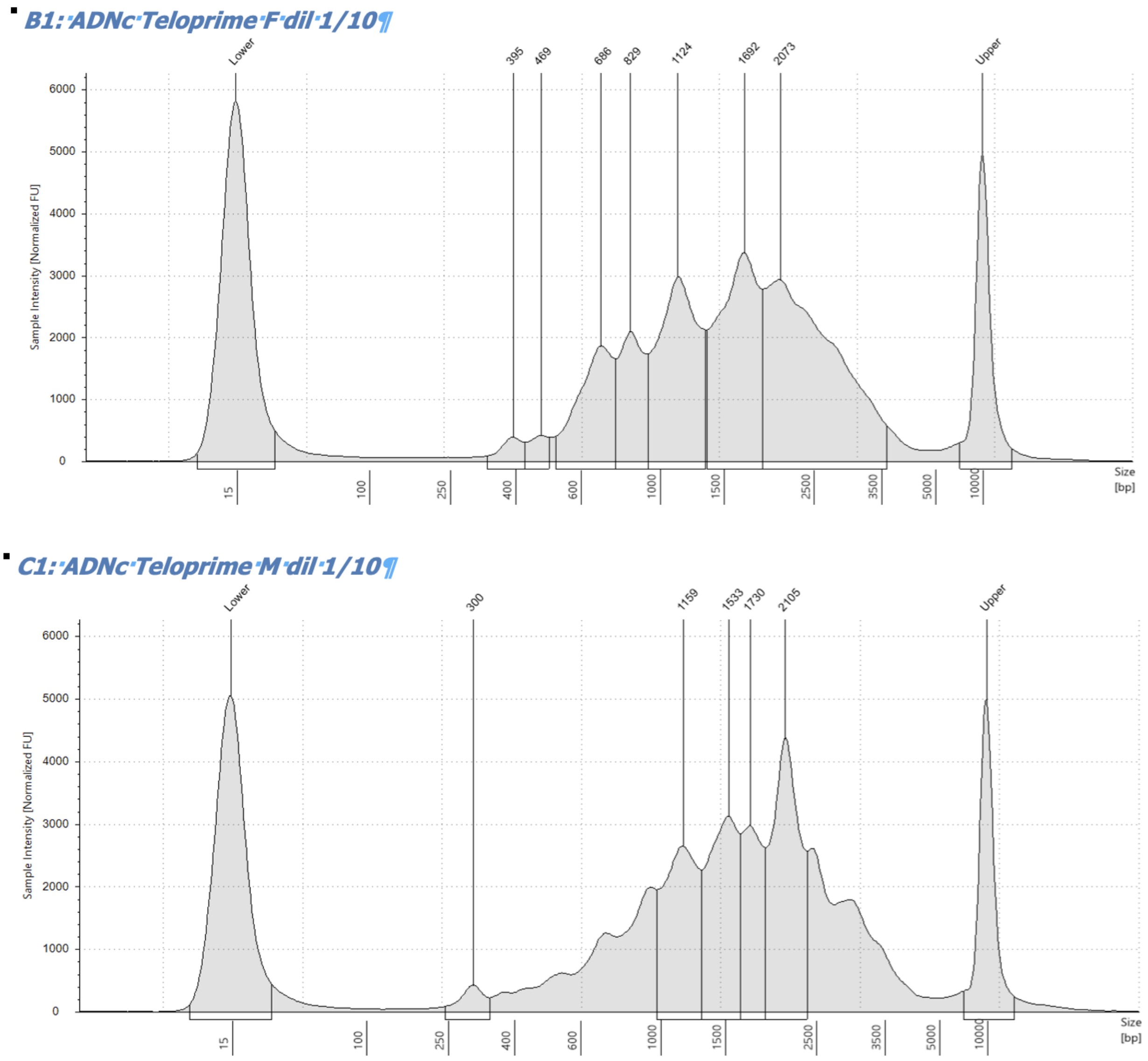


Figure S1. cDNA profile after TeloPrime amplification (F=> Female/ovaries; M=> Male/testes) showing cDNAs amplified are smaller than ~3.5 Kb. DNA ladder is shown as lower and upper peaks.

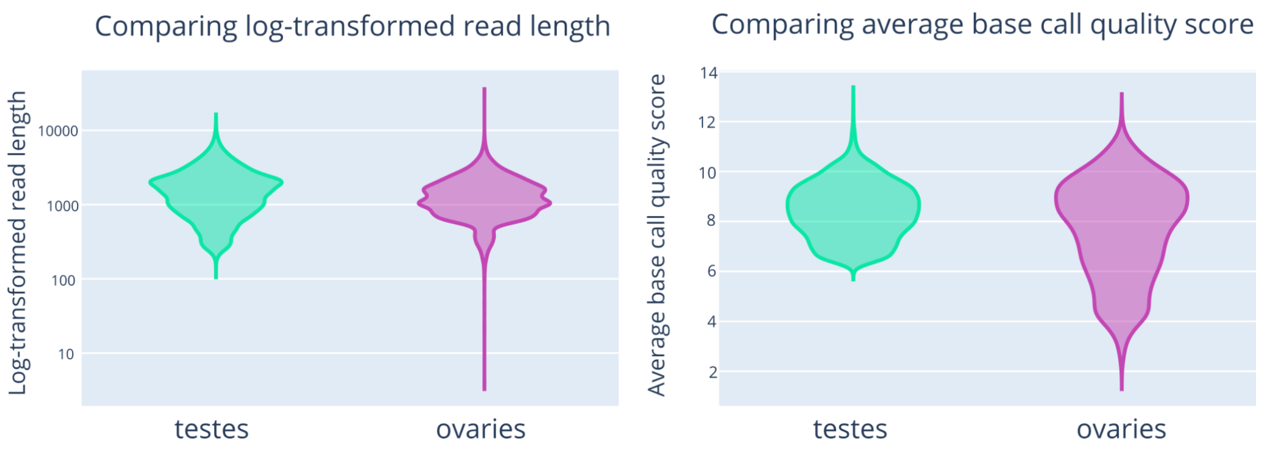


Figure S2. Read length and average base quality score for ONT reads obtained from ovary and testis cDNAs using the TeloPrime kit.

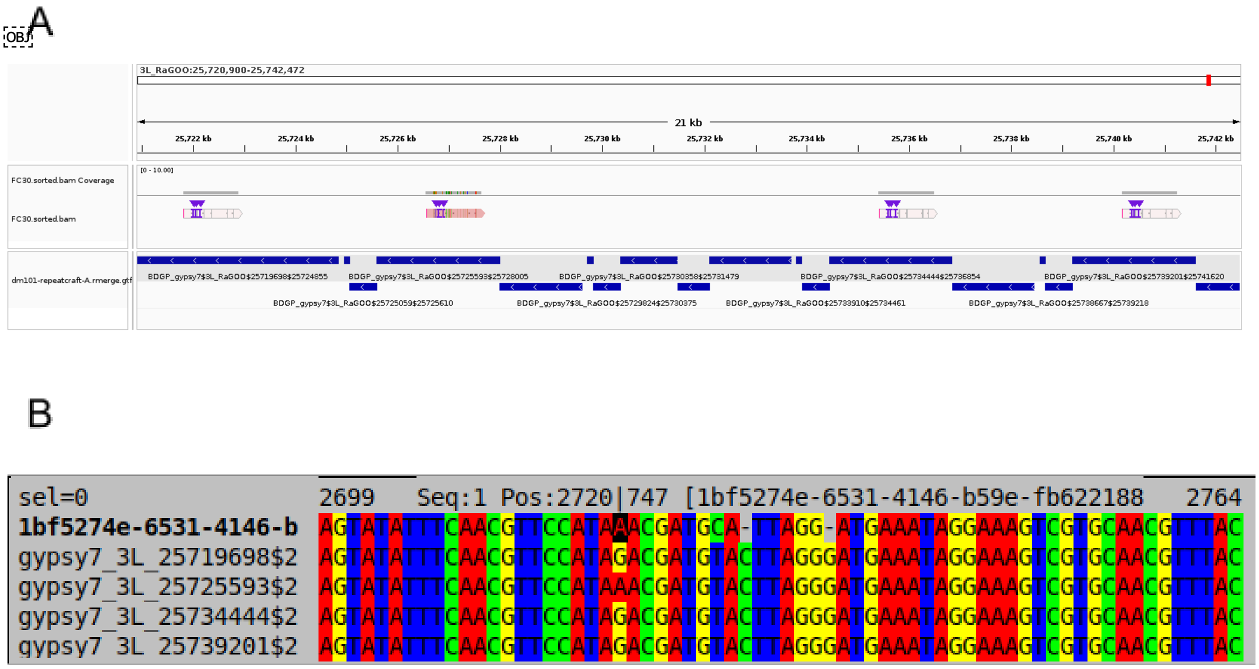


Figure S3. A. Example of a read mapping to four locations on the genome. These four locations are insertions of *BDGP_gypsy7*. The read aligns to these four locations with an alignment score of 861, 859, 859, 853 (mapping qualities 1, 0, 0, 0). The read is assigned to the location with the highest alignment score (AS tag in the sam format). B. Multiple sequence alignment of the read and the four insertions of *BDGP_gypsy7.* At position 2720 of the alignment, insertions 25719698, 25734444 and 25739201 contain a G, whereas insertion 25725593 contains an A. The read contains an A. Since this is the only site of divergence between the four copies, this alone explains the difference between alignment scores calculated by minimap2. Choosing the best score consists in assigning the read to insertion 25725593. The probability that the read stems from another insertion is 1/30 can be calculated as follows: the probability that there is a sequencing error at the site of divergence (1/10) and that the sequencing error yields an A and not a C or T (p=⅓). For cases of multi-mapping where there are n sites of divergence between the copies, assuming that the errors are independent, the probability of misassignment is (1/30)^n.

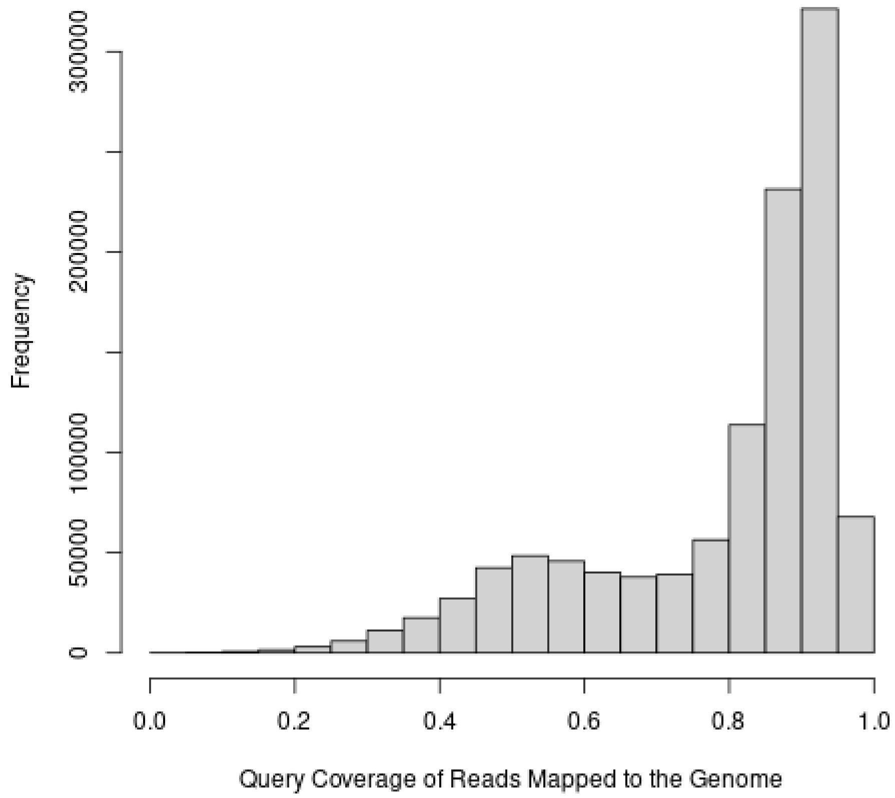


Figure S4. Histogram of the query coverage of reads mapped to the reference genome. 80% of reads have a query coverage centered on 90%, 20% of reads have a query coverage centered around 50%. Upon manual inspection, we find that the reads with a coverage centered around 50% stem from the sequencing of two independent RNAs that passed in the nanopore and were interpreted as a single molecule by the basecaller.

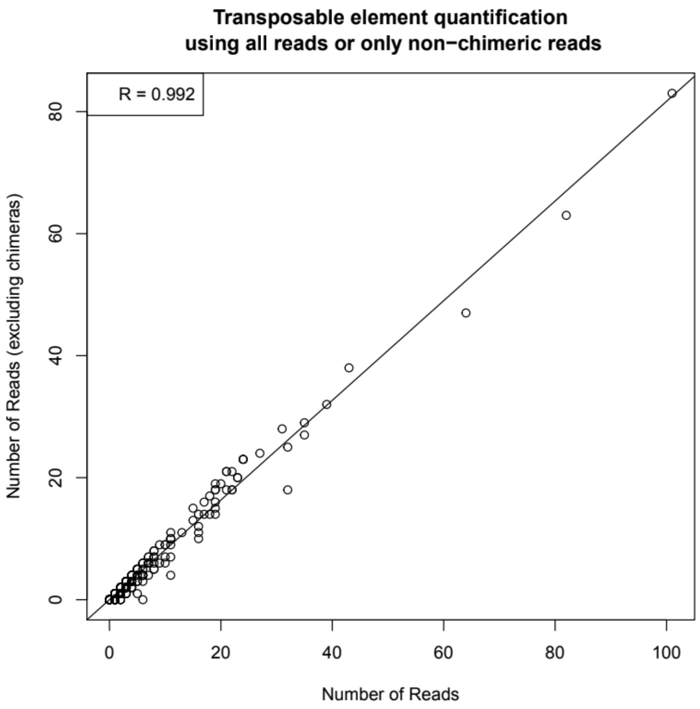


Figure S5. Transposable element quantification using all or non-chimeric reads only (defined in Figure S4 as reads spanning two cDNAs sequentially passing through the nanopore). Each dot corresponds to a TE insertion. The x-axis corresponds to the number of reads supporting the expression of this insertion when using all reads (including chimeras). In the y-axis, chimeric reads are excluded.

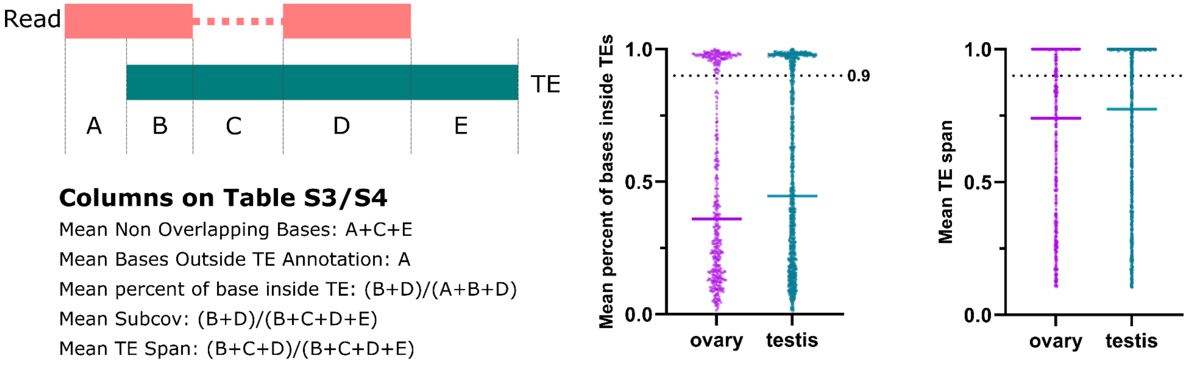


Figure S6. Left panel. Representation of the different metrics in Tables S3 and S4. Middle panel. Distribution of the metric “Mean percent of bases inside TEs” representing the average percentage of bases of a read within a TE sequence. The right panel depicts the distribution of the “Mean TE span” across the different TE copies.

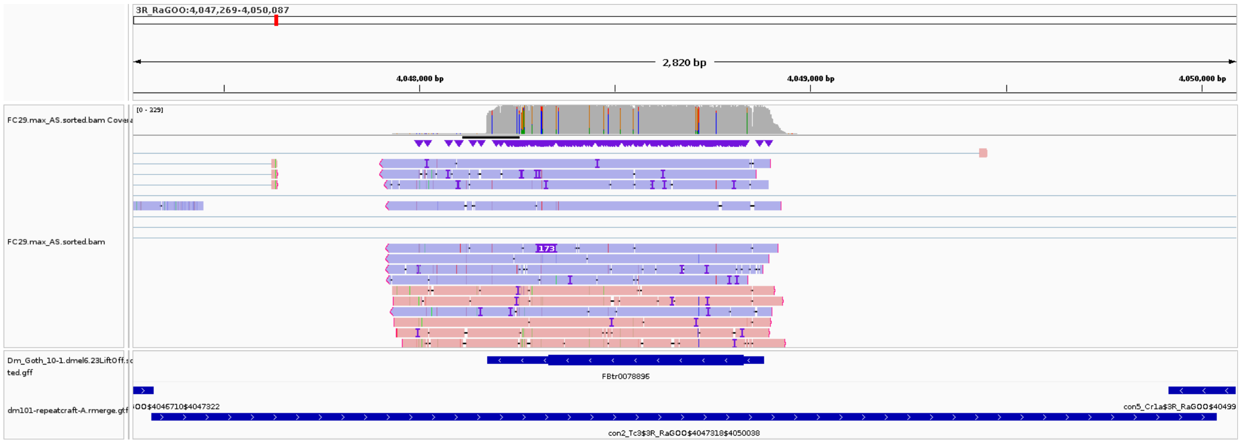


Figure S7. Example of a genomic region where both a gene and a TE are annotated. The reads map to both features, but the query coverage is much higher for the gene so the read is not assigned to the TE.

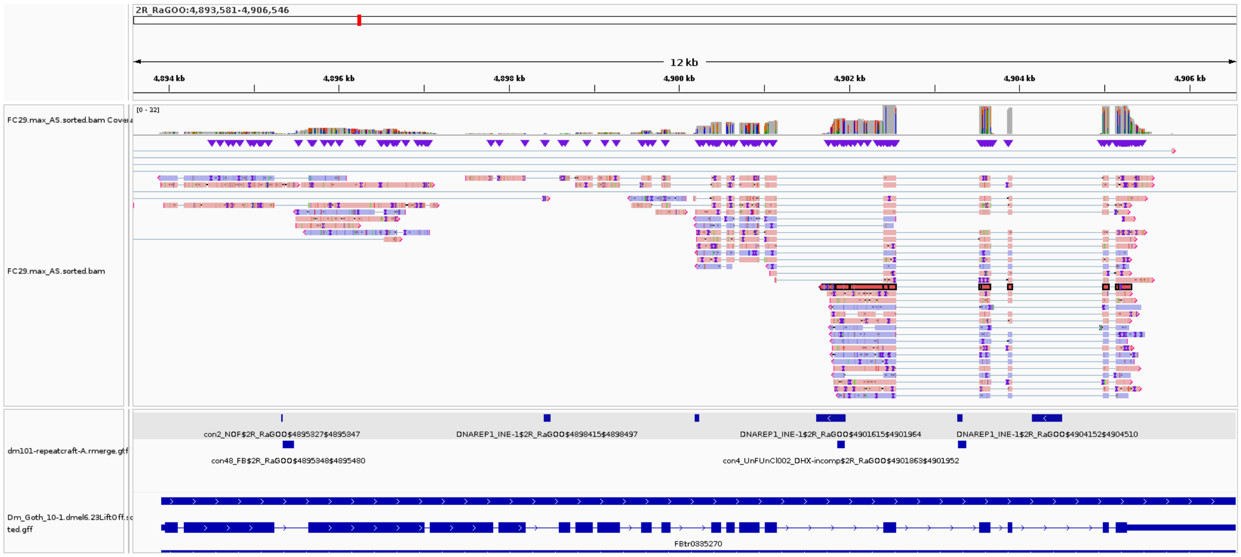


Figure S8. Example of a locus where reads overlap both a TE (DNAREP1_INE-1$2R_RaGOO$4901615$4901964) and a gene (*Gp210*, transcript FBtr0335270). The read highlighted in red is assigned to the TE because it covers better the TE than the gene. Reads located above this read in the track are assigned to the gene. Reads located below are assigned to the TE. We can hypothesize that this TE acts as an alternative promoter for this gene.

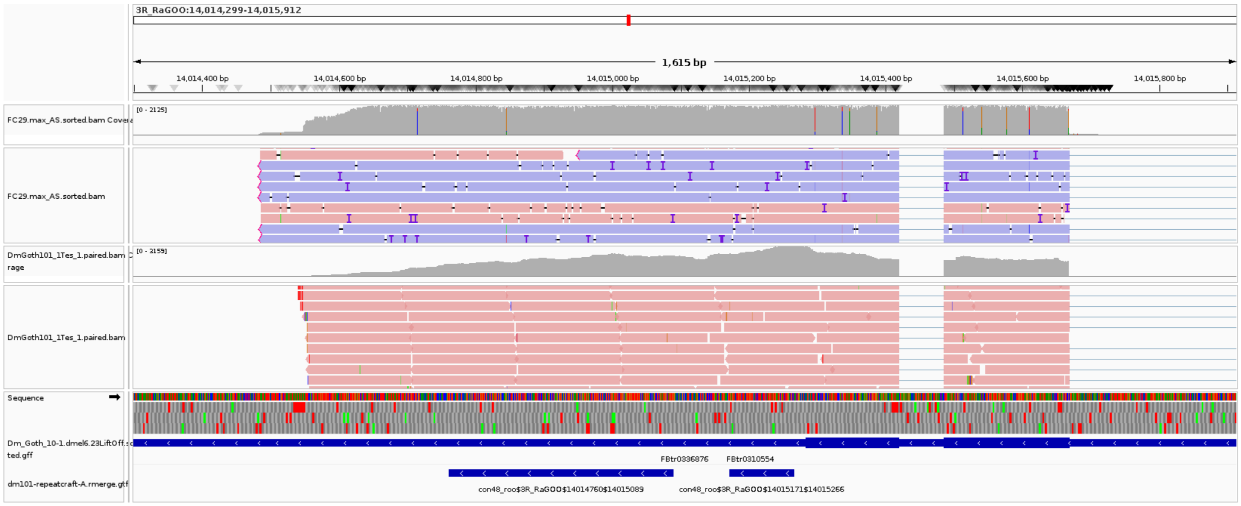


Figure S9: Mis-assignment of reads using short reads. Using short-reads, this *con48_roo* copy is estimated to be expressed (either by TETranscripts or Squire). Using long-reads, the reads are assigned to the gene, not the TE.

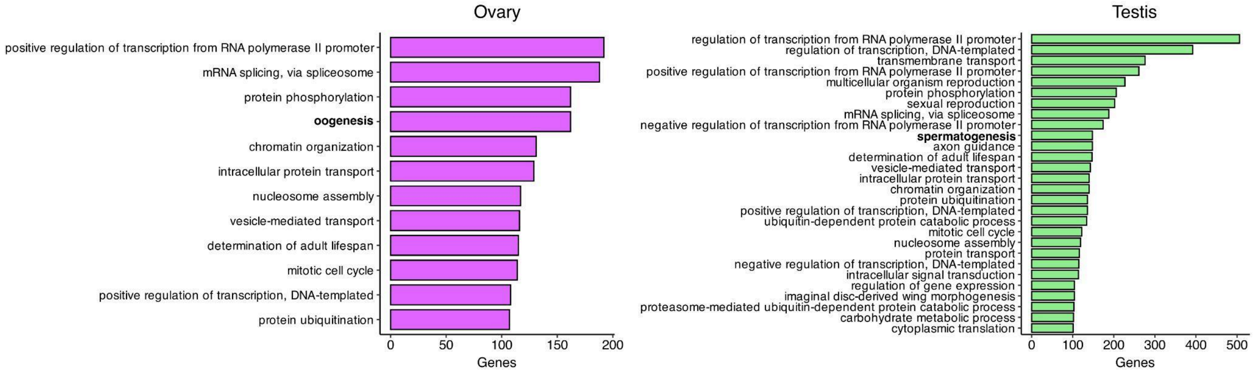


Figure S10. Gene Ontology enrichment analysis of long-read transcriptomes of ovaries and testis. In bold are enriched terms related to male and female gonadal development.

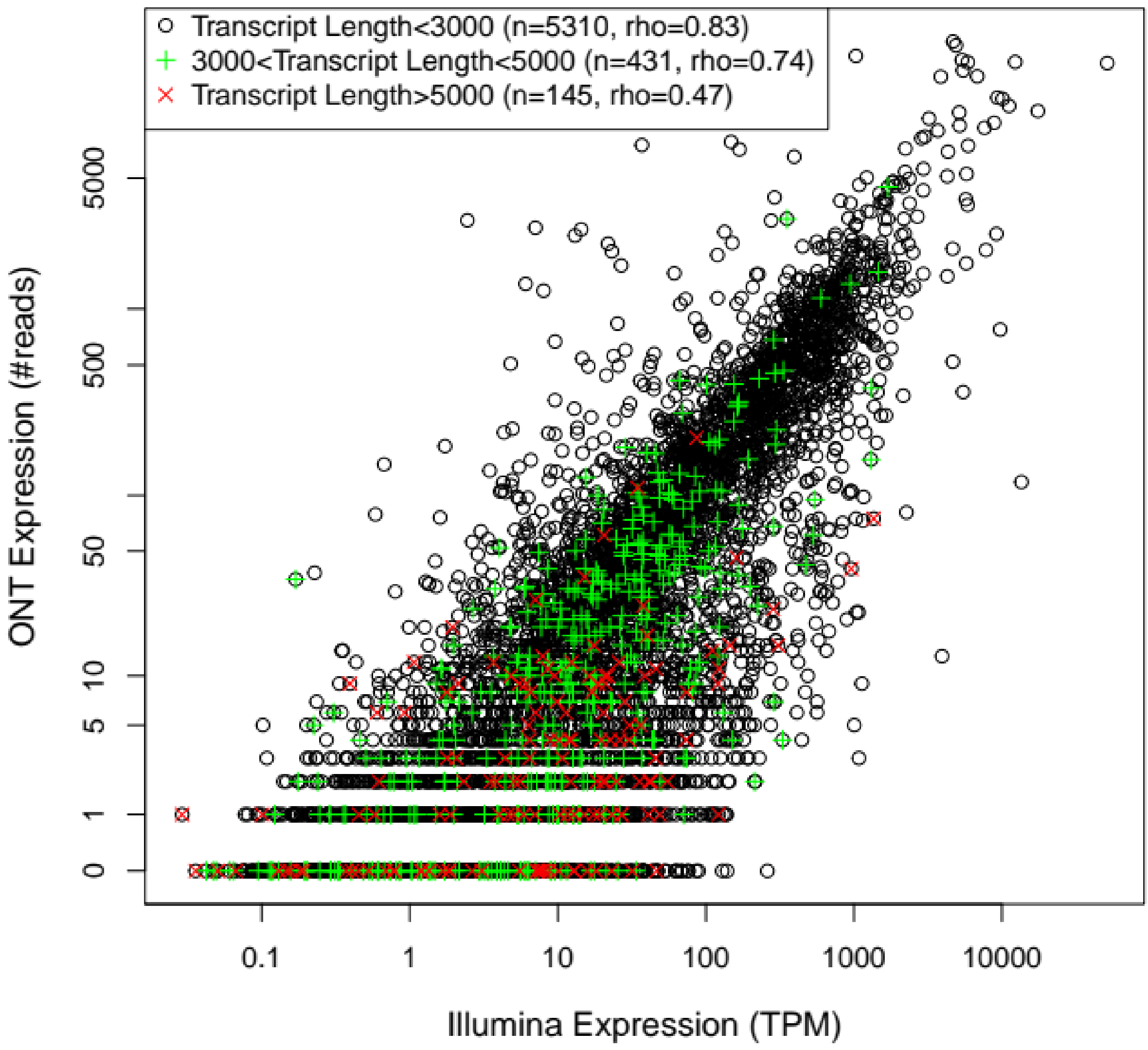


Figure S11: Gene expression quantification using Illumina and ONT sequencing in testes. Each dot is a gene with a single annotated isoform. Transcripts longer than 5 kb tend to be undersampled using TeloPrime.

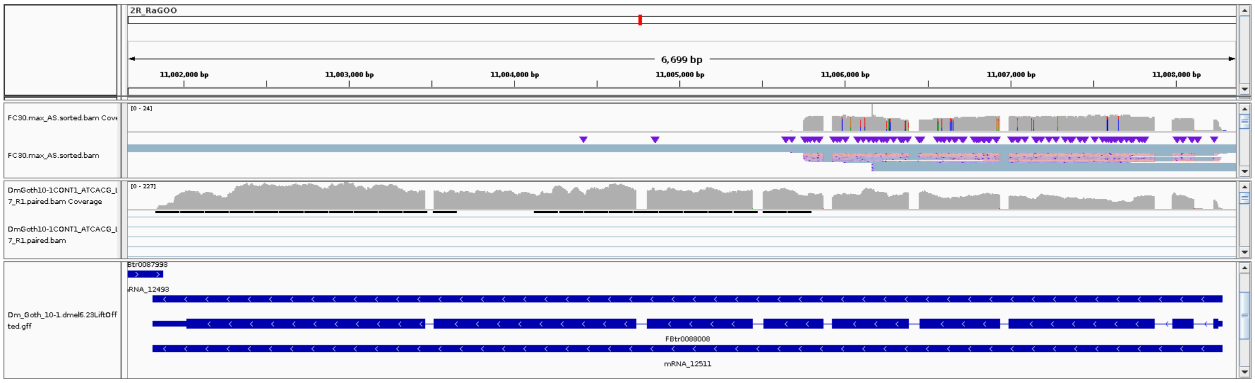


Figure S12: Example of a very long transcript, well captured by Illumina, but not by Teloprime Nanopore.

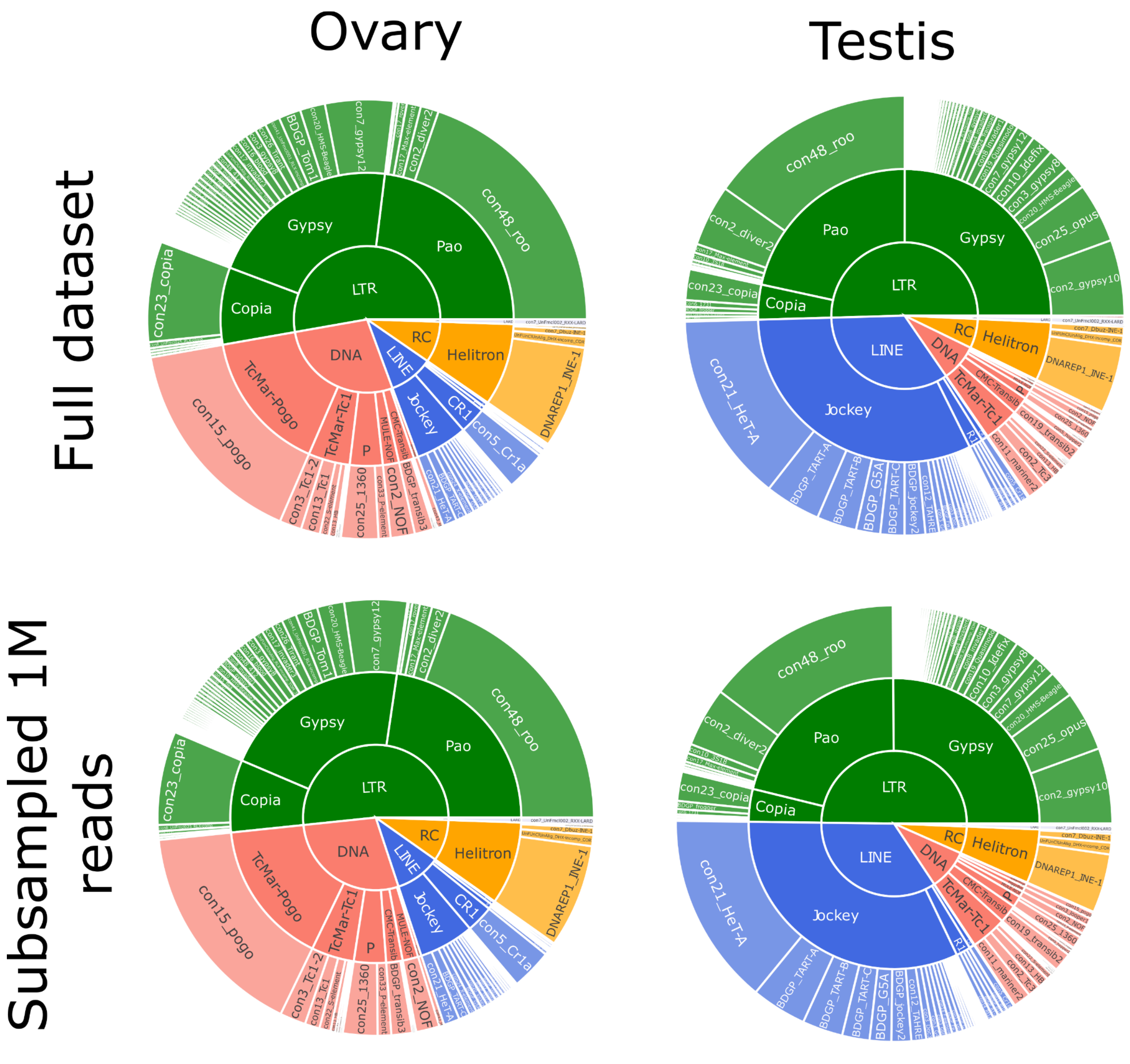


Figure S13. TE transcriptional landscape using the complete dataset as Figure 2B and a 1M subsampled ONT long-reads. The outer ring, middle ring and inside circle represent TE family, superfamily and subclass respectively. The area in the circle is proportional to the number of multi and uniquely mapped reads.

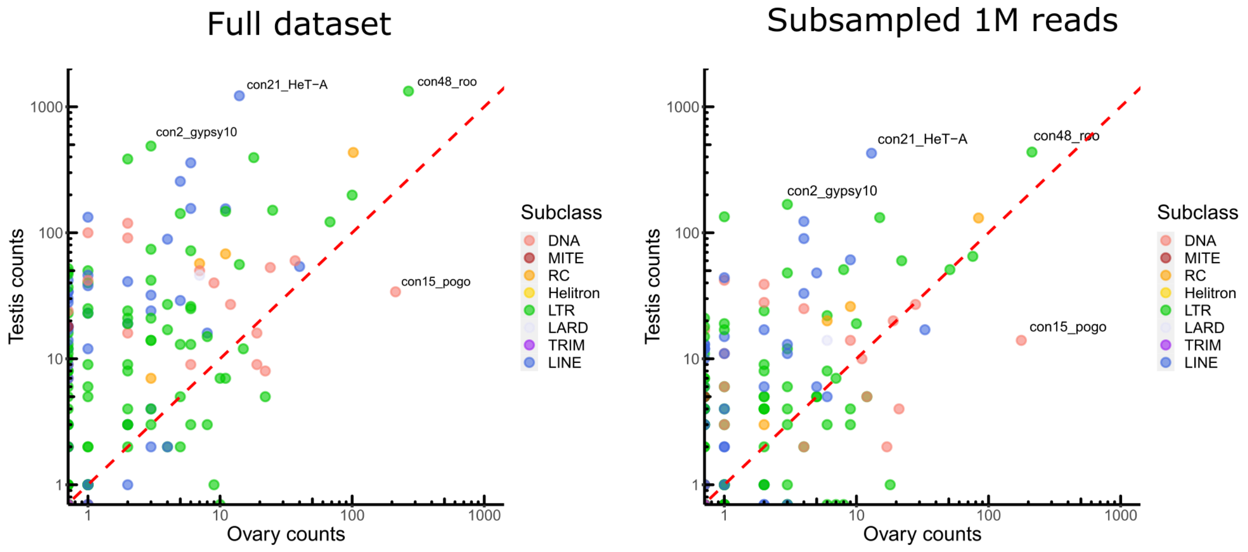


Figure S14. Comparison of TE expression ratio between testes and ovaries for the complete dataset as seen in Figure 2C and a 1M subsampled ONT long-reads.

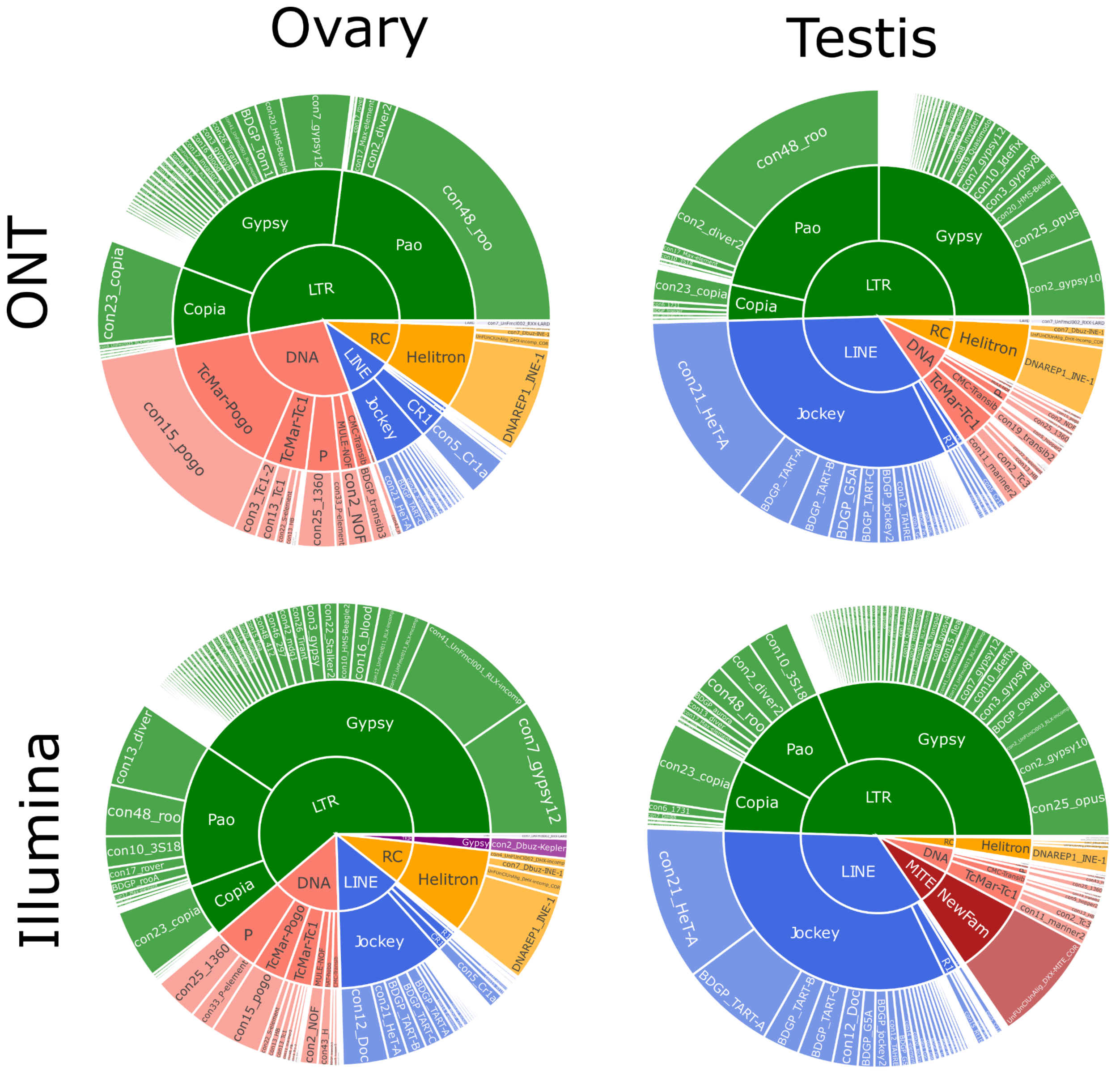


Figure S15. Global TE transcriptional landscape using ONT long-read sequencing (top panel, as Figure 2B) and Illumina short reads through TEtranscripts (bottom panel). The outer ring, middle ring and inside circle represent TE family, superfamily and subclass respectively. The area in the circle is proportional to the rate of expression.

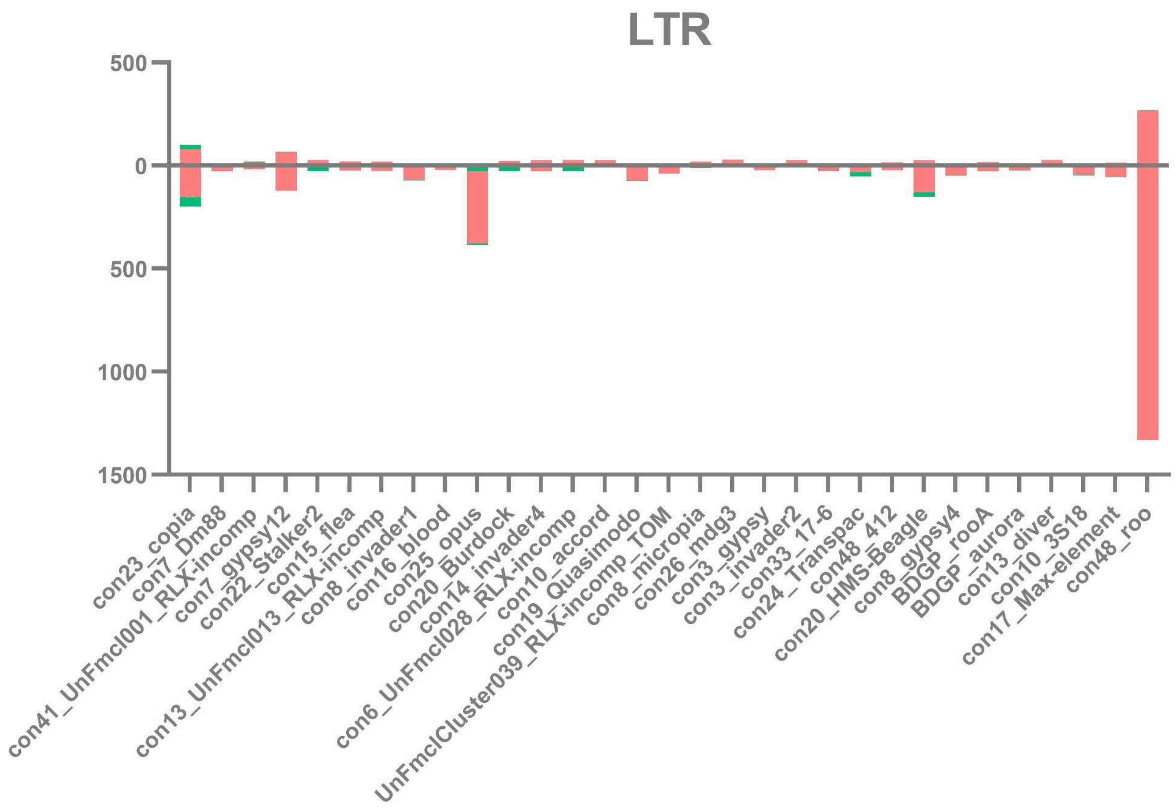


**Figure S16.** Multi-mapping and uniquely mapping ONT reads. **A.** Distribution of uniquely and multimapped reads across TE families in ovaries and testes (only TE families harboring at least one multimapped read are shown) for LTR elements, including *con48_roo* that has only one multimapping read in ovaries.

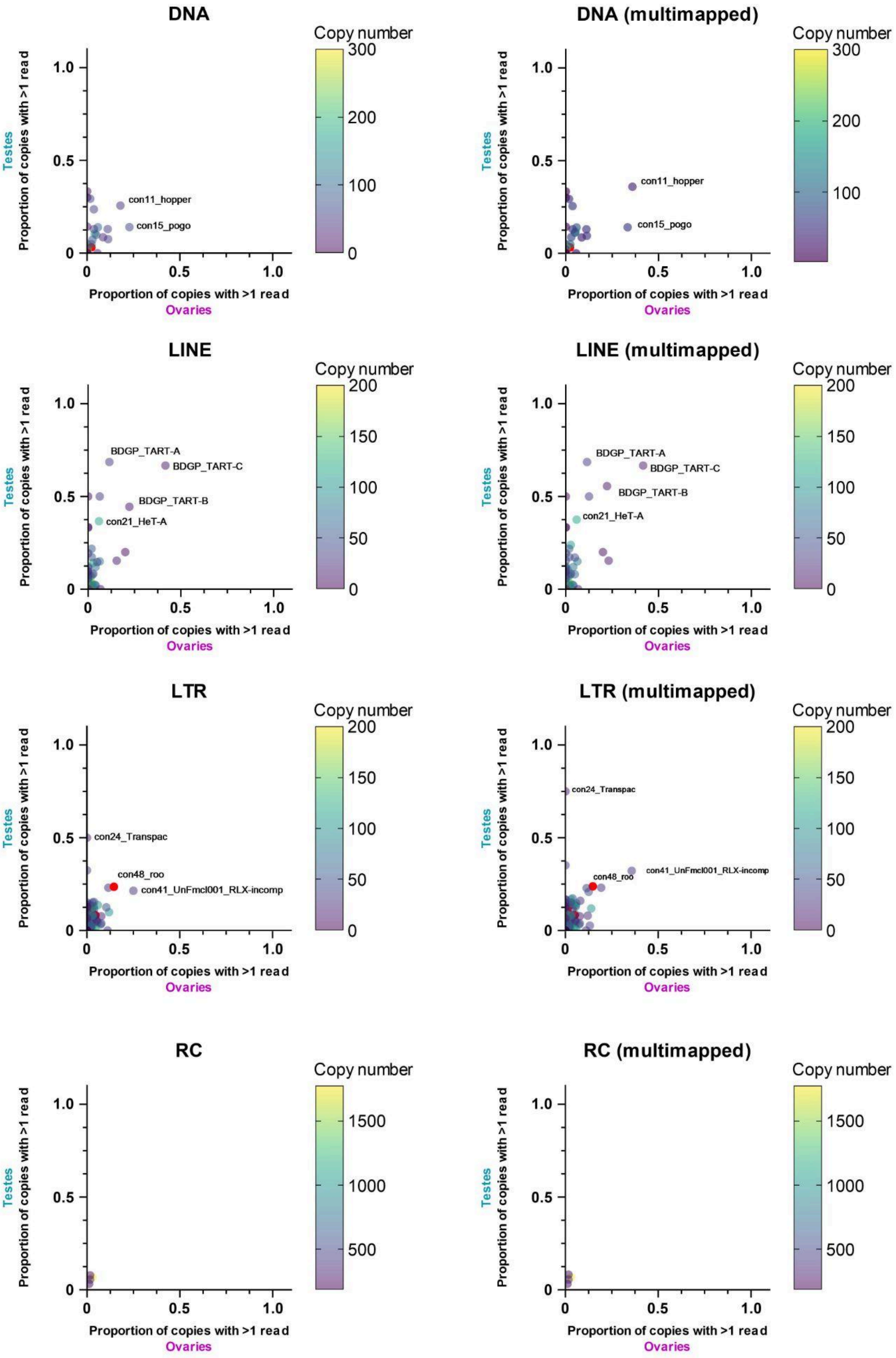
DNA
DNA (multimapped)
Copy number
con11_hopper
con15_pogo
LINE
LINE (multimapped)
BDGP_TART-A
BDGP_TART-C
BDGP_TART-B
con21_HeT-A
LTR
LTR (multimapped)
con24_Transpac
con48_roo
con41_UnFmcl001_RLX-incomp
RC
RC (multimapped)
Testes
Proportion of copies with >1 read
Ovaries

Figure S17. Frequency of transcribed copies (read > 1, uniquely and multi-mapped) within TE subfamilies in ovaries and testes, along with genomic copy number (color bar, 1 to 200 (LINE/LTR), 300 (DNA) copies) and 2000 for RC elements. All TE families harbouring more than 200 (LINE/LTR) or 300 (DNA) copies are depicted in pink. For DNA elements, *con25_1360* has 875 insertions. For LINE families, *con5_Cr1a* has 671 copies. The LTR families, *con10_idefix* (251), *UnFmclCluster039_RLX-incomp_COR* (278), *con19_Quasimodo* (253), *con2_diver2* (205), *con48_roo* (475), *con7_gypsy12* (242) and *con3_gypsy8* (315) are also depicted in pink.

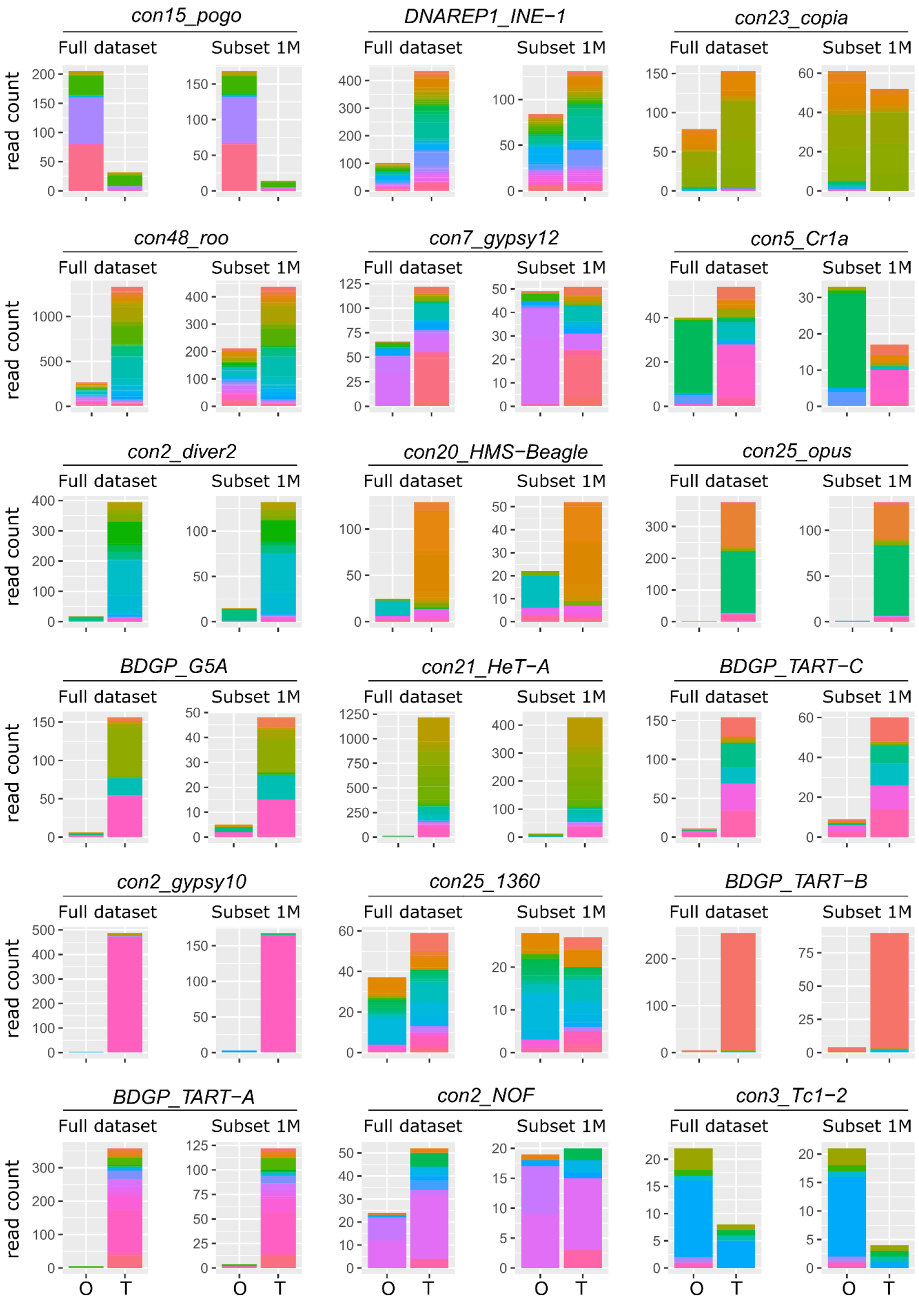


Figure S18. Distribution of read counts per copy for the 10 most expressed copies in ovaries and testes (18 TE families total), showing the overall expression of specific copies within a TE family (Table S3 and S4). For each TE family, both the complete dataset as seen in Figure 4B and the 1M subsampled dataset are shown. Copies are represented by different colors within the stacked bar graph. O: ovaries, T: testes.

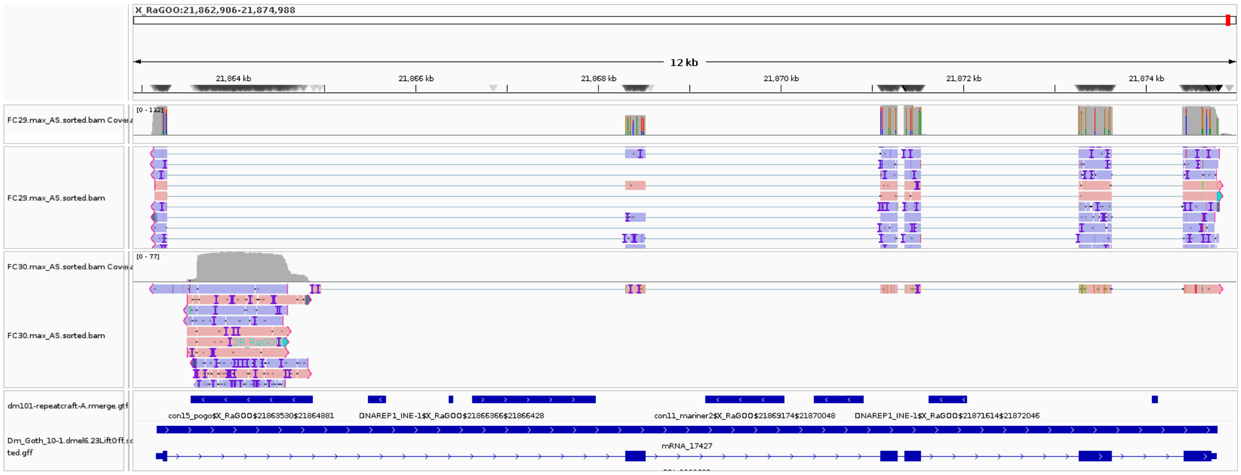


Figure S19. IGV screenshot of con15_pogo$X_RaGOO$21863530$21864881. Top track (FC29), testis coverage and an excerpt of mapped reads, followed by the same information for ovaries (FC30). Dmgoth101 repeat (dm101-repeatcraft-A.rmerge.gtf) and gene (Dm_Goth_10.1-dmel6.23LiftOFF) tracks are also shown and more information on the annotation can be seen in the material and methods section.

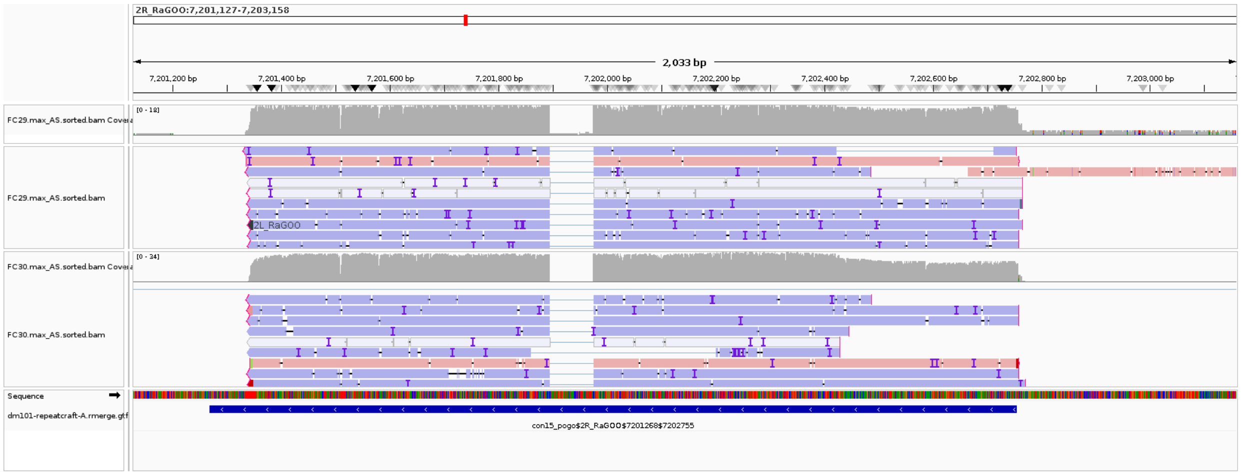


Figure S20. IGV screenshot of a *con15_pogo* copy. Top track (FC29), testis coverage and an excerpt of mapped reads, followed by the same information for ovaries (FC30). Dmgoth101 repeat (dm101-repeatcraft-A.rmerge.gtf) and gene (Dm_Goth_10.1-dmel6.23LiftOFF) tracks are also shown and more information on the annotation can be seen in the material and methods section.

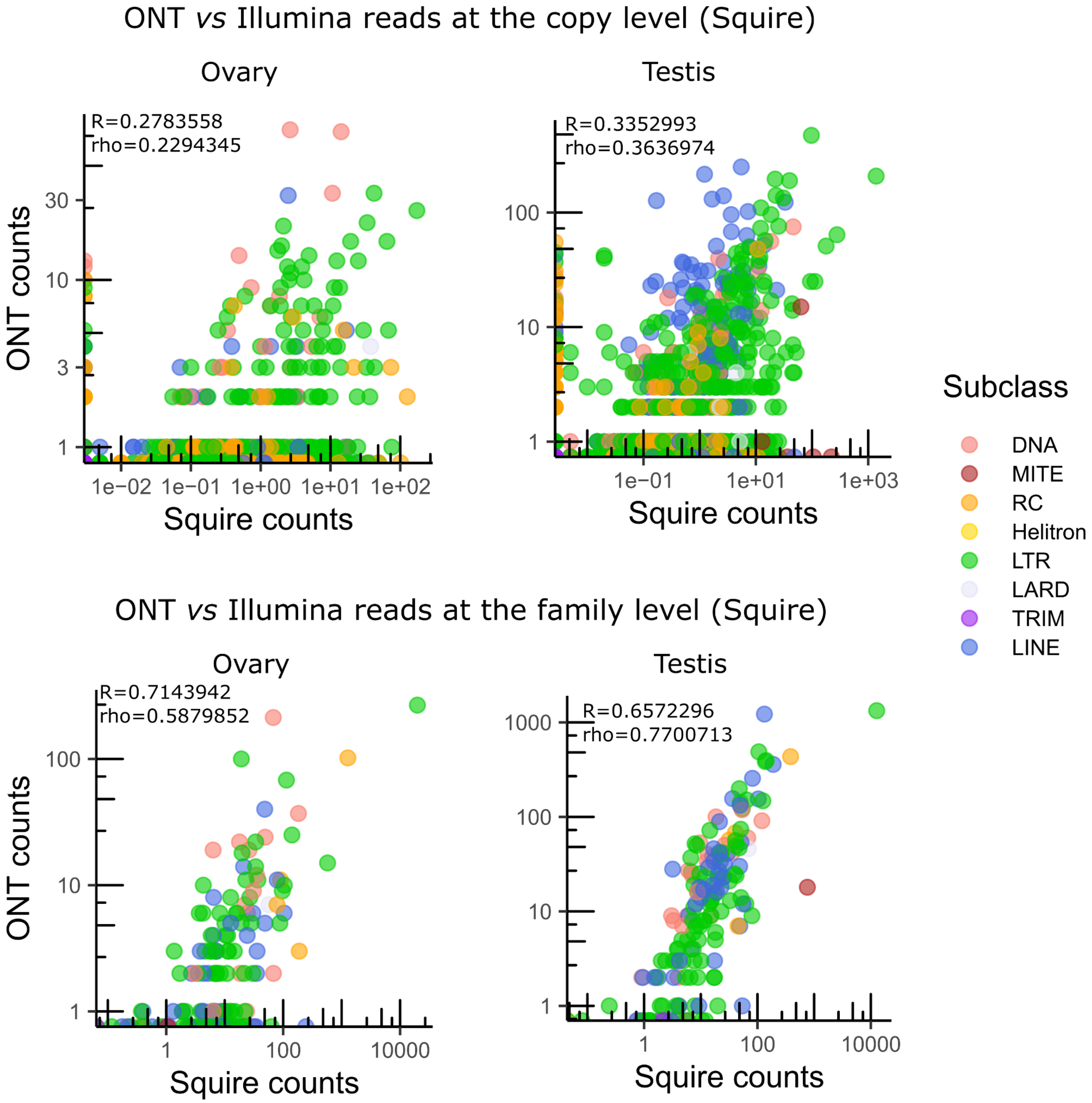


Figure S21. Comparison of ONT read count with Illumina short read on TE copy expression at the copy and family level. Short-read counts were performed with Squire.

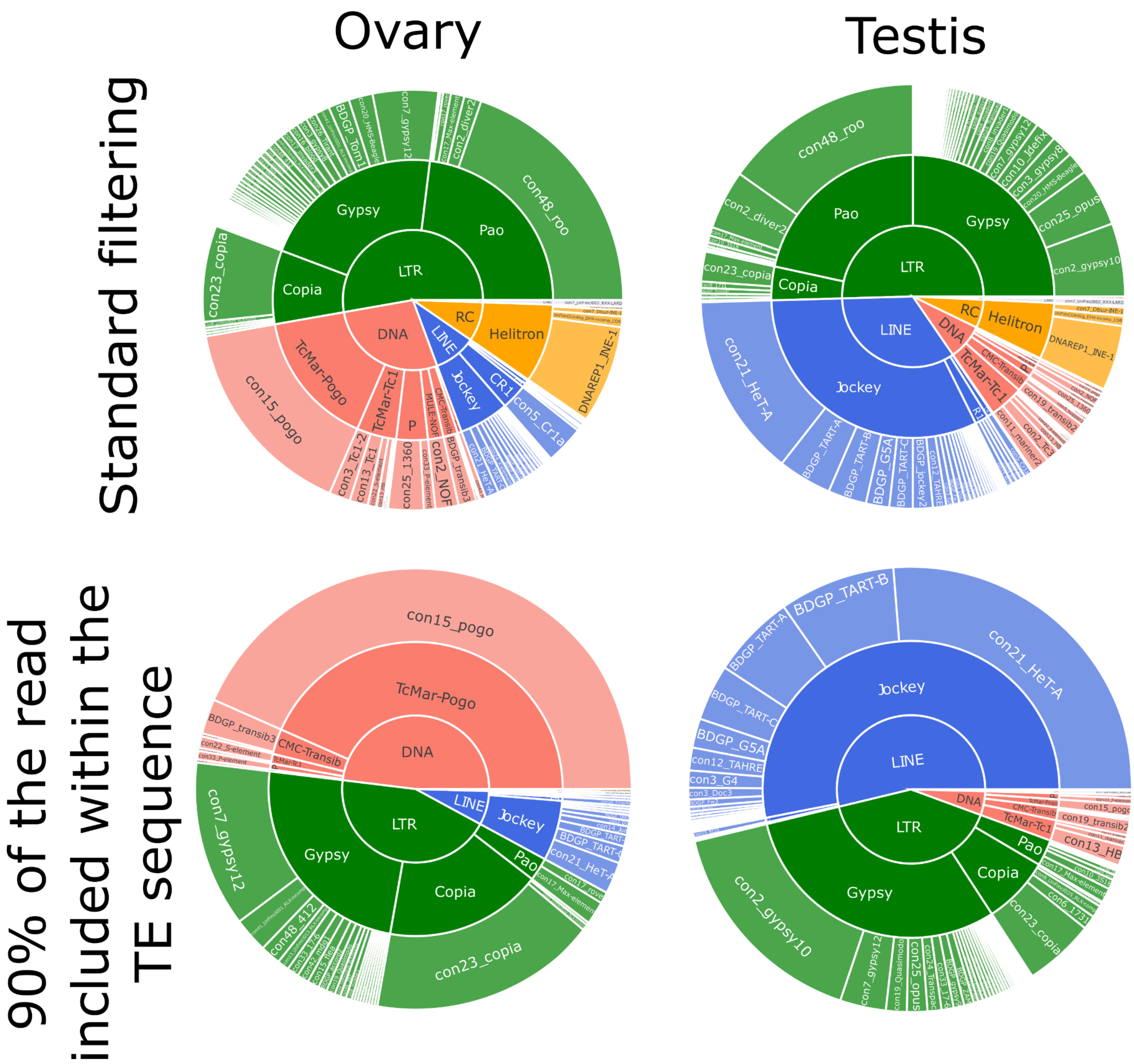


Figure S22. Global TE transcriptional landscape using ONT long-read sequencing with standard filtering (10% of the read sequence included within the TE) (as Figure 2B) and 90% of the read sequence within the TE (bottom panel). The outer ring, middle ring and inside circle represent TE family, superfamily and subclass respectively. The area in the circle is proportional to the rate of expression.

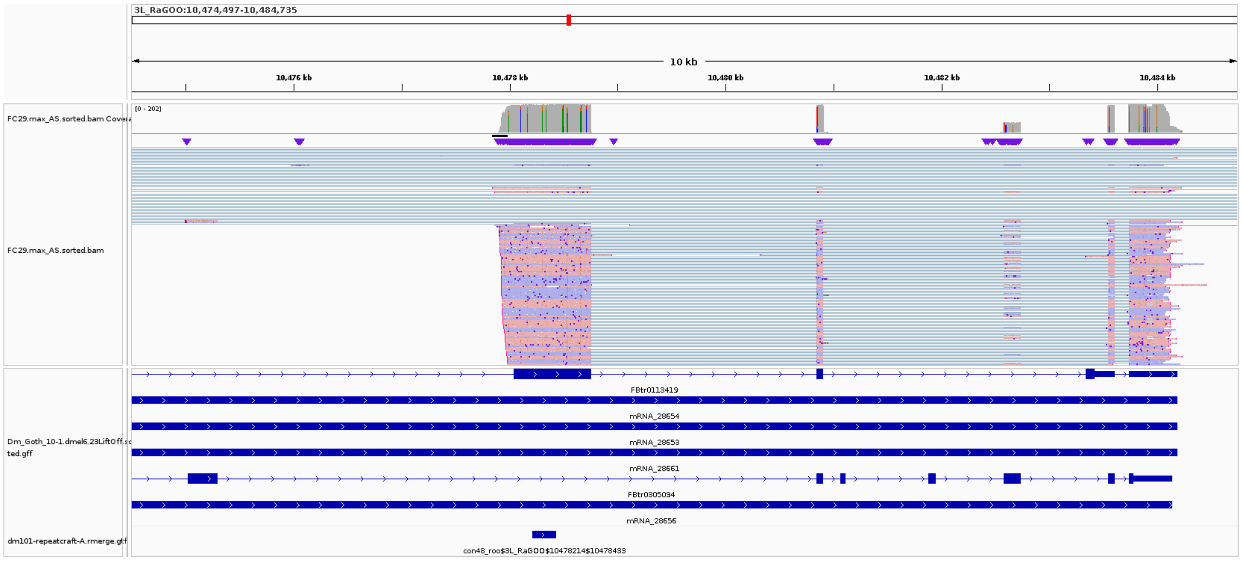


Figure S23. Example of *con48_roo* insertion where reads extend beyond the TE annotation. This insertion could act as an alternative promoter of its host gene.

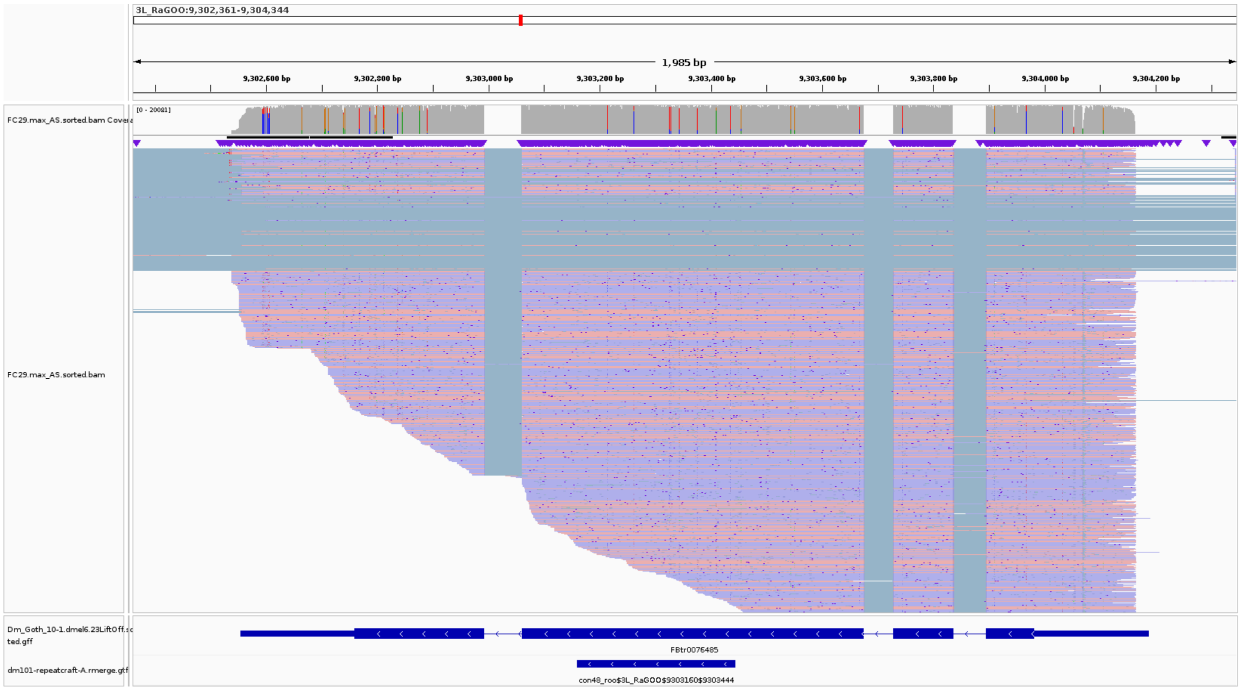


Figure S24. Example of a *con48_roo* insertion where reads extend beyond the TE annotation. This insertion is likely not active. It overlaps a highly expressed gene. Most reads are correctly assigned to the gene. However, because of the “staircase effect” (i.e. long reads do not always correspond to full-length transcripts), our algorithm wrongly assigns some reads to the TE, because these reads overlap more the TE than the gene.

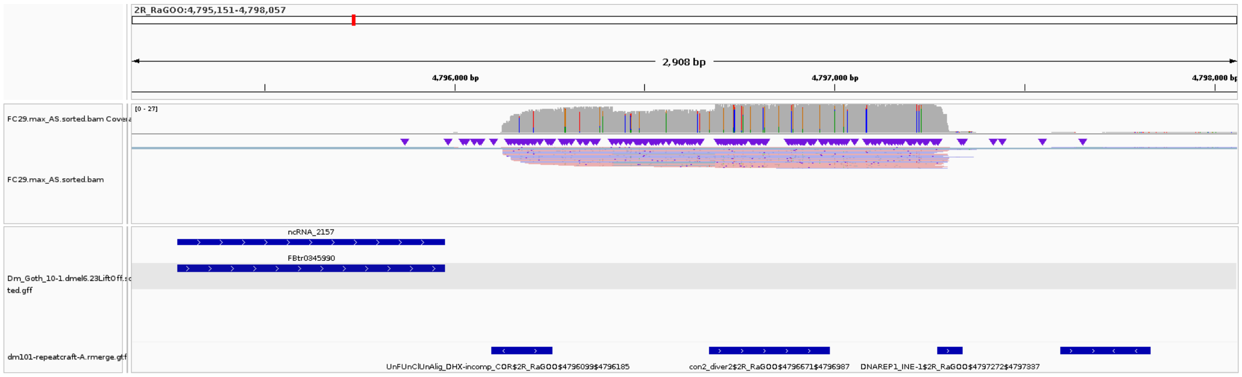


Figure S25. Example of a *con2_diver2* insertion where reads extend beyond the TE annotation. In this case, the transcriptional unit is not associated to a gene, but to three TE insertions (*con2_diver2, DNARep1_INE-1, UnfUnClUnAlig_DHX-incomp_COR*). Our algorithm assigns it to *con2_diver2*, which is the longest.

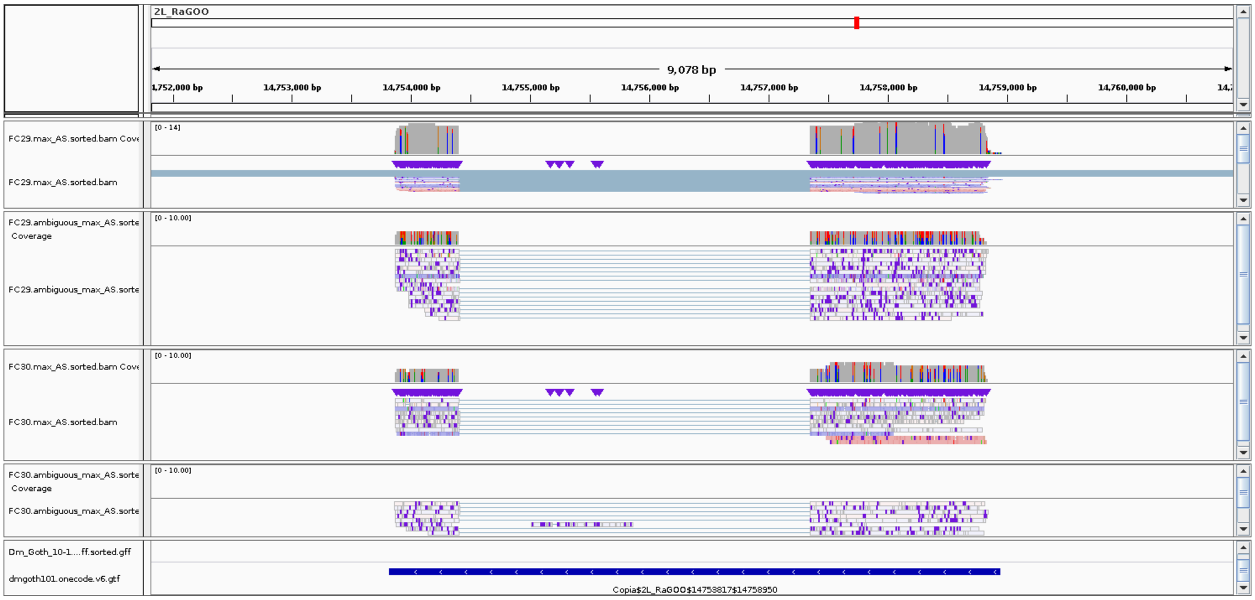


Figure S26. IGV screenshot of long reads mapping to Copia$2L_RaGOO$14753817$14758950). Track 1 and 3 are uniquely mapped reads (resp. in testes and ovaries). Track 2 and 4 are multimapped reads (resp. in testes and ovaries). Some reads in track 1 and 3 appear in white because those correspond to reads where the primary alignment was not the best alignment. This occurs in some cases because minimap's choice of primary alignment occurs early in the mapping algorithm and the choice may be incorrect. This assignment can be revised when looking at the alignment score.

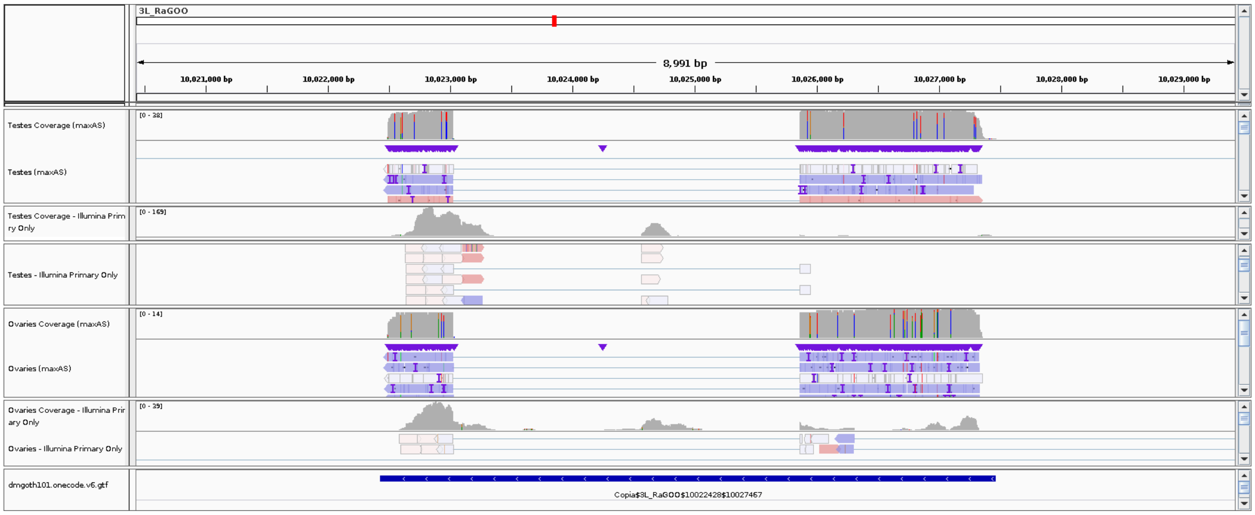


Figure S27. *con23_copia* splicing supported by uniquely-mapping reads. Top track, testis coverage and an excerpt of mapped reads, followed by the same information for short reads. Ovary tracks are depicted below. Dmgoth101 repeat and gene tracks are also shown and more information on the annotation can be seen in the material and methods section.

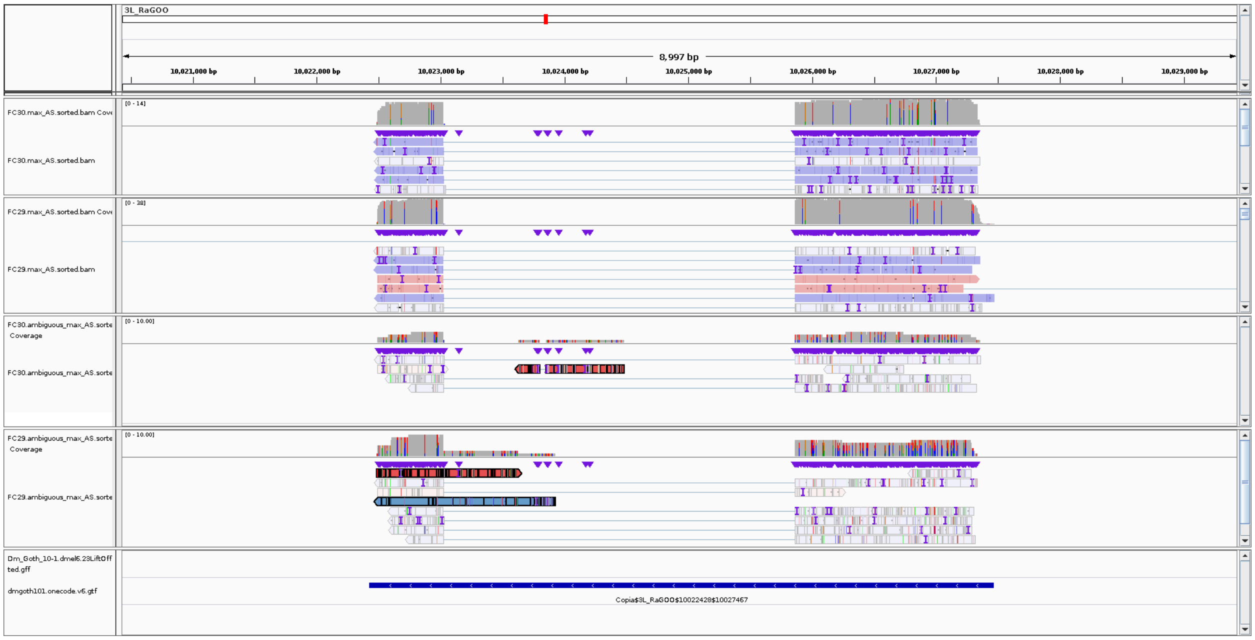

Figure S28: con23_copia$3L_RaGOO$10022428$10027467 long reads. Track 1 and 2 are uniquely mapped reads for ovaries and testes. Track 3 and 4 are multimapped reads for ovaries and testes. Highlighted read map in the intron of *con23_copia*, suggesting that the unspliced version of *con23_copia* could be present in the sample, but not correctly sequenced, possibly due to uncompleted reverse transcription.

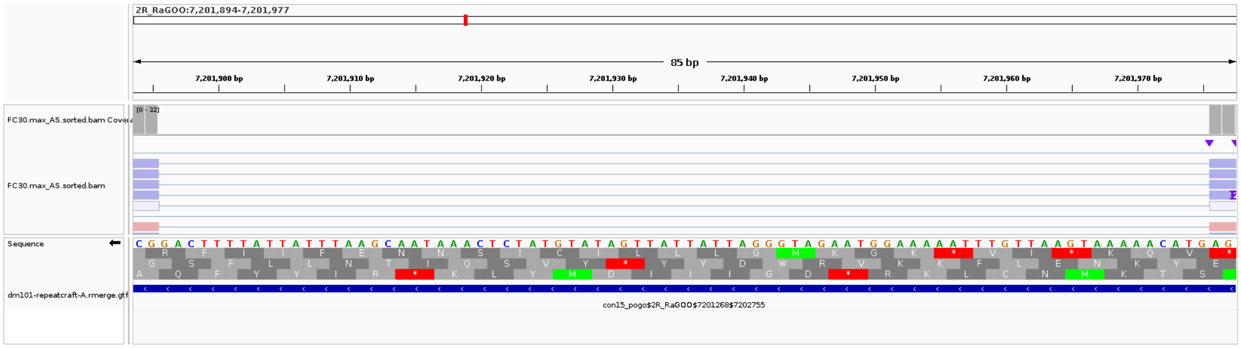

Figure S29: Zoom on the intron of con15_pogo$2R_RaGOO$7201268$72027559. The TE is annotated on the minus strand. The donor site is GT. The acceptor site is AG.

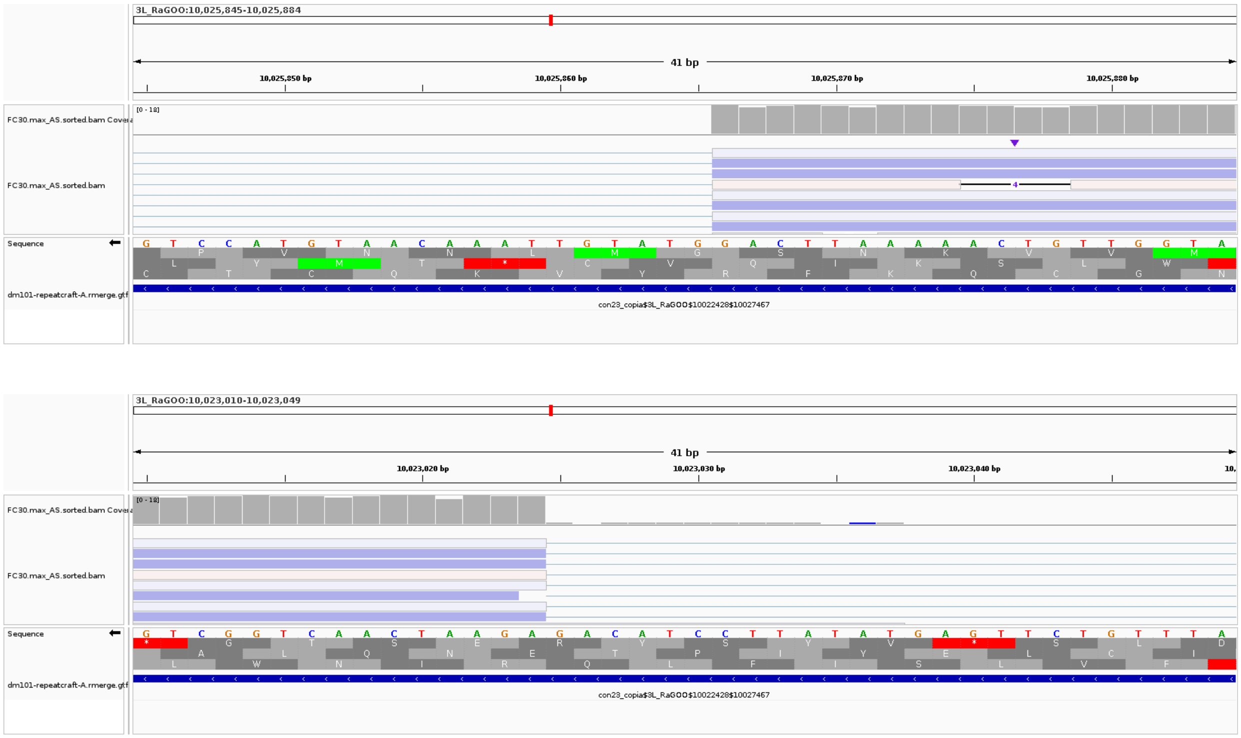


Figure S30: Zoom on the donor site and acceptor site of the intron of con23_copia$3L_RaGOO$10022428$10027467. The TE is annotated in the minus strand. The intron is 2.8kb long.

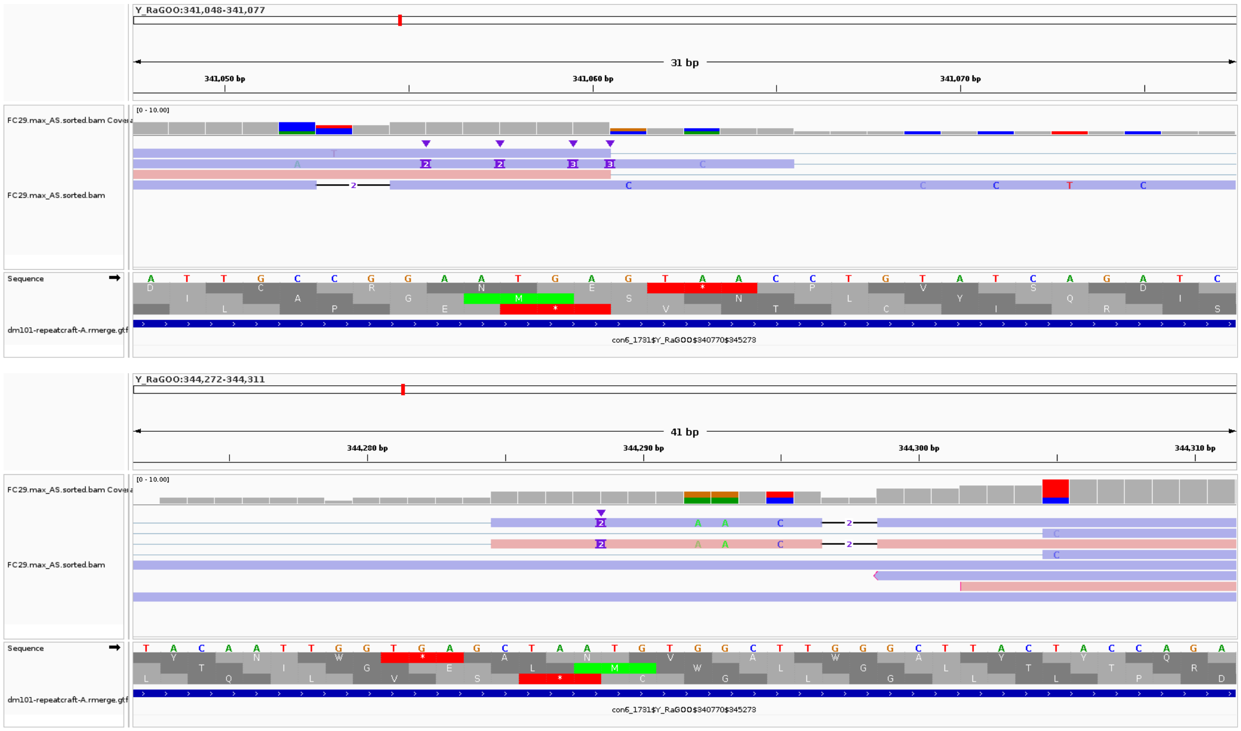


Figure S31: Zoom on the donor site and acceptor site of the intron of con6_1731$Y_RaGOO$340770$345273, two reads (the 1st and the 3rd) are spliced using a GT-AG consensus.

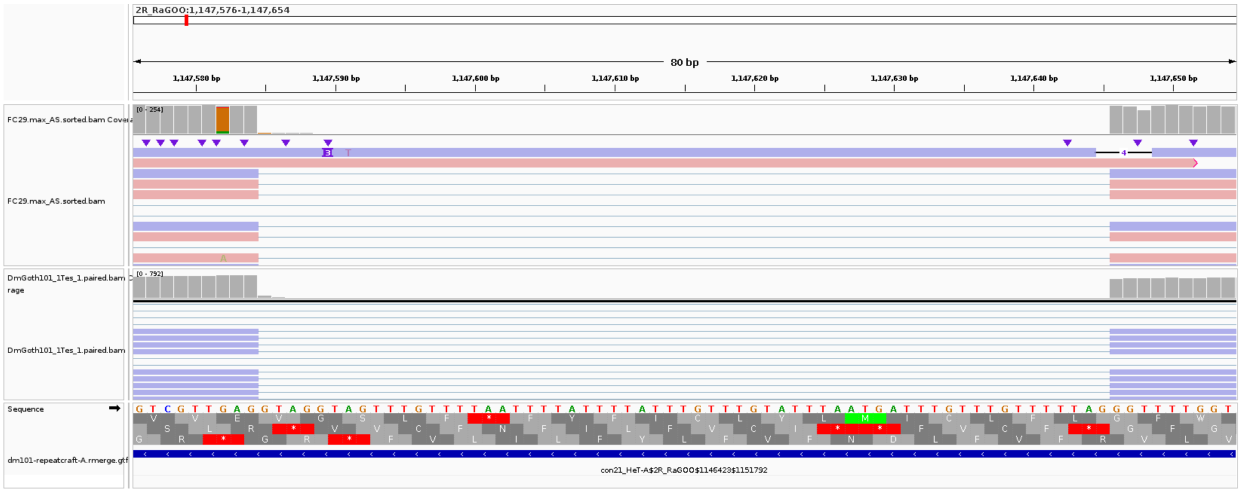


Figure S32: Zoom on the intron of con21_HeT-A$2R_RaGOO$1146423$1151792. The TE is annotated on the minus strand (arrows pointing to the left in the Repeat Track). The intron is CT-AC on the minus strand. It is GT-AG on the plus strand (the sequence shown is the plus strand, arrow pointing to the right). The stranded short reads all map on the plus strand (blue colour), supporting the hypothesis that the transcript is antisense.

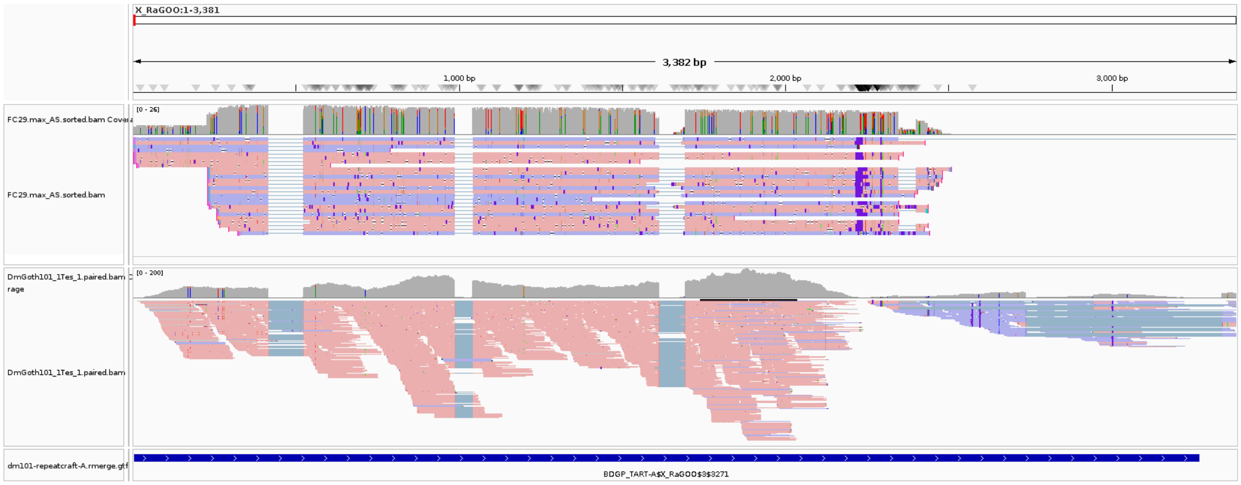


Figure S33: Insertion BGDP_TART-A$X_RaGOO$3$3271 is located at the telomere of the X chromosome. Long reads exhibit gapped alignments. Gaps are flanked by CT-AC consensus. The insertion is located on the direct strand of DNA. Stranded short reads all map to the reverse strand (pink colour) which supports the hypothesis that the transcript is antisense.

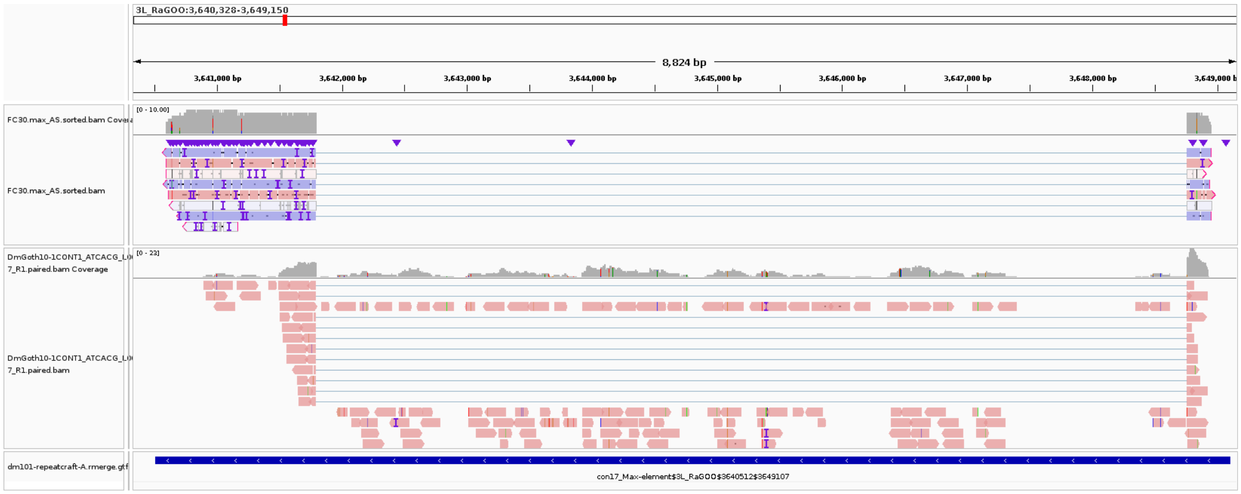


Figure S34: Full-length genomic copy con17_Max-element$3L_RaGOO$3640512$3649107 is detected as spliced using long reads, while both the spliced and unspliced version are detected using short reads. The unspliced version is very long (>8kb) and may be more difficult to detect using long reads. Additionally, it is poorly expressed.

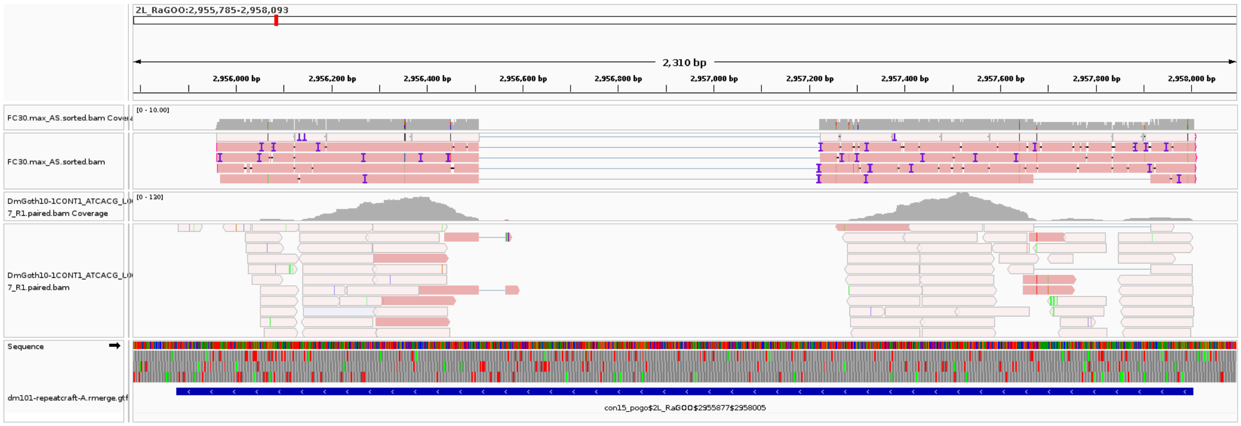


Figure S35: Full-length genomic copy of *con15_pogo*, con15_pogo$2L_RaGOO$2955877$2958005 is detected as spliced using long reads. Short reads do not support the unspliced version. No short read seems to map to the exon junction either. This could be due to the difficulty to map short reads to novel unannotated exon junctions, in particular in repeated regions.

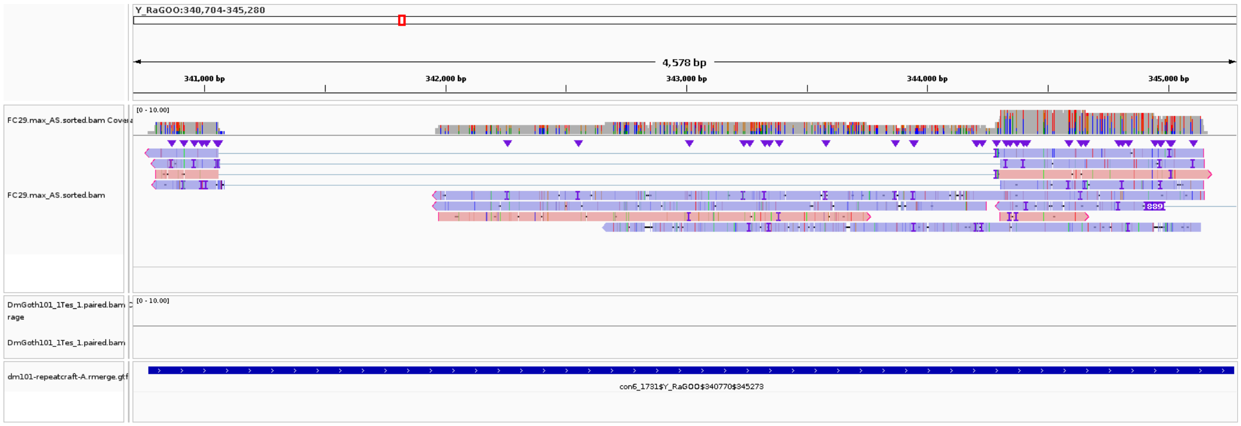


Figure S36: Full-length genomic copy of *con6_1731*, con6_1731$Y_RaGOO$340770$345273 is detected as spliced using long reads. It is not detected at all using short reads. This could be due to its very low expression level.
